# Early postnatal DNA methylation dynamics define neuronal subtypes and are disrupted by MECP2 loss

**DOI:** 10.64898/2026.06.06.730504

**Authors:** Lauren E. Rylaarsdam, Ruth V. Nichols, Brendan L. O’Connell, Sabrina Kragness, John F. Yung, Arpiar Saunders, Gail Mandel, Andrew C. Adey

## Abstract

DNA methylation is the foundational layer of epigenetic regulation underlying mammalian tissue specification. During synaptogenesis, neurons accumulate high levels of a unique type of methylation in which a cytosine precedes a C, A, or T (mCH) instead of the canonical guanine (mCG). Disruption of the mCH reader methyl-CpG binding protein 2 (MECP2) causes the devastating neurodevelopmental disorder Rett syndrome, yet the role of mCH in neurodevelopment remains unclear. In this study, we generated the first large-scale single-cell methylation atlas of the early postnatal mouse brain and resolved subtype-specific trajectories of canonical and noncanonical methylation changes underlying neuronal subtype specification. Each identified population undergoes a rapid maturation event of the noncanonical methylome occurring between the first and second postnatal weeks, with highly subtype-specific changes concentrated at genes involved in synaptic partner establishment. Trajectory analysis resolved that subtype diversification, such as the separation of parvalbumin and somatostatin-positive GABAergic interneurons, is further facilitated by a hierarchical sequence of methylation changes in genes involved in regulation of membrane potential. In parallel, we applied single-cell methylation analysis to a mouse model of Rett syndrome that recapitulates severe human symptoms. We show that while noncanonical methylation accumulation proceeds in each subtype in the absence of MECP2, populations that have been consistently implicated in Rett pathology such as GABAergic interneurons have an increasing number of differentially methylated regions with age and fail to accumulate typical global levels of mCH. Together, our study defines the methylation dynamics that facilitate neuronal subtype specification and resolves subtype-specific, global disruptions of noncanonical methylation in Rett syndrome.

## Main Text

Neurodevelopment is an intricate phenomenon that requires careful orchestration of both extrinsic and intrinsic processes. A small pool of progenitors ultimately give rise to the entire brain, differentiating into thousands of distinct populations using the same genetic material.(*1*) Intrinsically, the first layer of epigenetic control mediating interactions between DNA and the environment is the deposition and removal of methyl groups to the 5’ position of cytosines, known as DNA methylation.(*2*) Methylation is a critical mechanism for fine-tuning gene expression via the recruitment or inhibition of transcription factors to the DNA.(*2*) Even subtle deviations in methylation patterns can have devastating consequences, highlighted by the relevance of proteins with epigenetic-modifying functions to disorders such as cancer, autism, intellectual disability, epilepsy, and schizophrenia.(*3–12*)

In mammalian tissues, methylation is canonically deposited by DNA methyltransferases (DNMTs)(*13*, *14*) at cytosines that precede guanines (mCG). This dinucleotide repeat is enriched at regulatory regions like promoters and enhancers.(*15*, *16*) Hypermethylation will canonically lead to gene silencing(*17*, *18*) through inhibition of transcriptional machinery or recruitment of repressive factors to the DNA. Though it is a relatively stable epigenetic mark, mCG can be removed via an oxidative process initiated by ten-eleven translocases (TETs), which convert methylcytosine to hydroxy-methylcytosine (hmC).(*19–22*) In this manner, methylation facilitates modulation of gene expression to meet the needs of the cell. DNA methylation is most dynamic during embryonic development, when an initial wave of demethylation characterizes cells departing from a pluripotent state, followed by widespread re-methylation as tissues are specified.(*23–25*) In adulthood, the mCG landscape continues to be remodeled more subtly in response to age and other environmental factors.

Many decades after the discovery of mCG, it was established that neurons - a particularly long-lived and specialized class of cells - further leverage a distinct noncanonical methylation context(*26–29*) hypothesized to be an added layer of epigenetic regulatory control.(*30*) At birth, the brain is like other mammalian tissues with the majority of CG sites methylated genome-wide.(*27*, *31*) As the brain begins to mediate interactions with the environment, an explosion of synaptogenesis and circuit refinement occurs, coinciding with the DNTM3A-mediated methylation of millions of cytosines preceding non-guanine nucleotides (A, C, T; or mCH).(*27*, *32*) The resulting neuronal mCH levels are so abundant that total mCH sites may exceed mCG.(*27*, *33–35*) Astrocytes and oligodendrocytes also accumulate mCH at about five-fold lower levels, but not microglia, which arise from the endodermal lineage.(*27*) mCH levels fluctuate across broad megabase-scale topologically associated domain (TAD) boundaries.(*27*, *34*, *36*) Within TADs, mCH levels are further sub-bounded by gene bodies - particularly at critical long neuronal genes - where hypermethylation is associated with transcriptional repression.(*30*, *32*, *34*, *36–38*) The resulting noncanonical methylation patterns are the most subtype-specific epigenetic modality that has been described.(*27*, *33–35*, *39*, *40*) Loss of mCH organizer NSD1 results in impaired establishment and maintenance of neuronal identities,(*41*) further supporting the framework of mCH as a critical layer of epigenetic regulatory control that facilitates delineation of closely related neuronal subtypes.(*30*, *32*, *36*, *37*, *42*) However, knowledge of how mCH is accumulated has either been inferred from bulk bisulfite sequencing at early postnatal stages or bulk and single-cell snapshots in the adult,(*27*, *33–35*, *38*) leaving outstanding questions as to the role of noncanonical methylation in facilitating neuronal specification and why the brain needs to maintain it at such high levels.

In addition to the timing of its accumulation, the importance of mCH to neuronal function is further highlighted by the devastating consequences upon loss of its only known reader, methyl-CpG binding protein 2 (MECP2),(*43*) which results in Rett syndrome.(*44–46*) Patients meet initial milestones but start progressively deteriorating within the first few years of life. Symptoms include loss of ability to coordinate purposeful movement, repetitive hand motions, severe intellectual disability, seizures, breathing issues, and a myriad of other debilitating problems.(*47*, *48*) Nearly all patients are female(*47*, *48*) as *MECP2* lies on the X chromosome(*44*) and complete loss is ultimately lethal, but rare cases also exist of male patients with somatic mosaicism.(*49–52*) While MECP2 also binds mCG, an abundance of evidence points to mCH being critical to Rett pathology, including: symptom onset occurs just following the mCH accumulation plateau;(*27*) mice with a MECP2 methyl binding domain that cannot bind to mCH - but can still bind mCG - exhibit severe Rett-like phenotypes;(*53*) and genes that accumulate elevated mCH after birth are preferentially mis-regulated in mouse models.(*43*) Finally, the distribution of methylation across contexts within the mCH classification mirrors MECP2 binding affinity, with the mCAC trinucleotide sequences showing the strongest enrichment.(*42*)

Extensive research efforts to uncover the mechanisms of Rett syndrome have revealed that MECP2 reaches histone-octamer levels(*54*) during the early postnatal period to bind ubiquitously across the genome(*43*) and perform highly cell type-specific, activity-dependent functions throughout life.(*55–68*) MECP2 is canonically understood to act as a transcriptional repressor, particularly at long neuronal genes in highly methylated TADs.(*37*, *42*, *69–72*) However, there are nuances to this, as MECP2 also binds strongly to a subset of enhancer regions independent of methylation status(*73*, *74*) and can function as an activator.(*75*, *76*) Additionally, MECP2 plays widespread roles in epigenetic processes including chromatin organization,(*54*, *75*, *77–79*) the formation of TADs,(*36*) and may directly interact with the methyltransferase DNMT3A.(*80*) Systematic ablation of MECP2 across distinct lineages using genetic strategies in mice have pinpointed the contributions of various cell types to Rett symptoms: for example, selective loss of MECP2 in parvalbumin-positive interneurons causes motor, sensory, memory, and social deficits;(*81*) loss in somatostatin-positive interneurons leads to seizures and stereotypies;(*81*) loss in excitatory neurons causes cortical hyperexcitability;(*64*) and loss in the brainstem contributes to breathing difficulties.(*62*) The distinct cell type-specific pathologies could partially explain why perturbances from bulk tissue studies appear widespread and subtle,(*82*) highlighting the need for single cell-based strategies to further understand the mechanistic etiology of Rett syndrome.

Here we leveraged sciMETv3,(*83–85*) a high-throughput single-cell combinatorial indexing approach for methylation analysis, to investigate the accumulation of non-CG methylation in the developing brain. We generated a comprehensive single-cell methylation atlas of the early postnatal mouse brain across four time points spanning the critical period of noncanonical methylation accumulation, revealing the hierarchical epigenetic changes that accompany the emergence of diverse neuronal subtypes. Across subtypes, we identified a critical inflection point to an adult-like methylome state between the first and second postnatal weeks - a period coinciding with critical events like eye opening(*86*) and the GABAergic excitatory to inhibitory switch.(*87*, *88*) Within subtypes, we identified that genes involved in synaptic partner establishment and regulation of membrane potential had the most dynamic methylation state.

Furthermore, to better understand the role of non-CG methylation in cortical development and the epigenetic landscape of Rett syndrome, we extended our analysis to mice with a c.G311A variant in *Mecp2*. This patient-derived truncating variant (p.W104X) eliminates the DNA binding domain of MECP2,(*89*, *90*) causing severe motor and respiratory symptoms that are recapitulated in the mouse model.(*90*) This model was originally generated because the G311A mutation is therapeutically tractable through ADAR2-mediated RNA editing, a strategy that avoids pathological MECP2 overexpression(*46*) and has successfully rescued respiratory phenotypes *in vivo*.(*90–92*) Our single-cell methylation analysis in MECP2^W104X/y^ neurons revealed selective populations, including parvalbumin and vasopressin-positive GABAergic interneurons, increasingly accumulate differentially methylated regions with age and fail to accumulate normal global levels of mCH upon MECP2 loss. This is the first evidence of a shift in the noncanonical methylation landscape in Rett syndrome, which has not been resolved in previous studies using bulk-cell approaches. Altogether, results provide new insight into the trajectory of methylation changes facilitating neuronal specification and challenge prior understandings of the epigenetic landscape of Rett syndrome.

### A single-cell atlas of DNA methylation in the postnatal mouse brain reveals rapid maturation across subtypes between the first and second postnatal weeks

We first sought to characterize the hierarchy of methylation changes defining specification towards the adult state to yield insight into the function of non-CG methylation in the developing brain. Previous studies have pinpointed postnatal weeks one to four as a key period of epigenomic restructuring in mice,(*27*, *38*, *93*) but the sequence of methylation events within this timeframe had never been resolved at the granular, subtype level. We therefore microdissected the murine rostral and caudal cortex, hippocampus, striatum, and thalamus at four early postnatal time points spanning the vast majority of mCH accumulation (P7, P14, P21, P28).(*27*) Neurons, which accumulate five-fold greater mCH levels than glia, were enriched by sorting for NeuN^+^ nuclei.(*24*, *27*, *33*) Methylomes were then isolated in high-throughput by leveraging a combinatorial indexing-based strategy for methylation analysis (sciMETv3; (*83*, *84*, *94*) **Fig. 1A**). Altogether, we profiled 163,524 passing single-cell methylomes across twelve mice spanning four time points and five brain regions (**Fig. 1B-E**; see **Table S1**).

**Figure 1.**
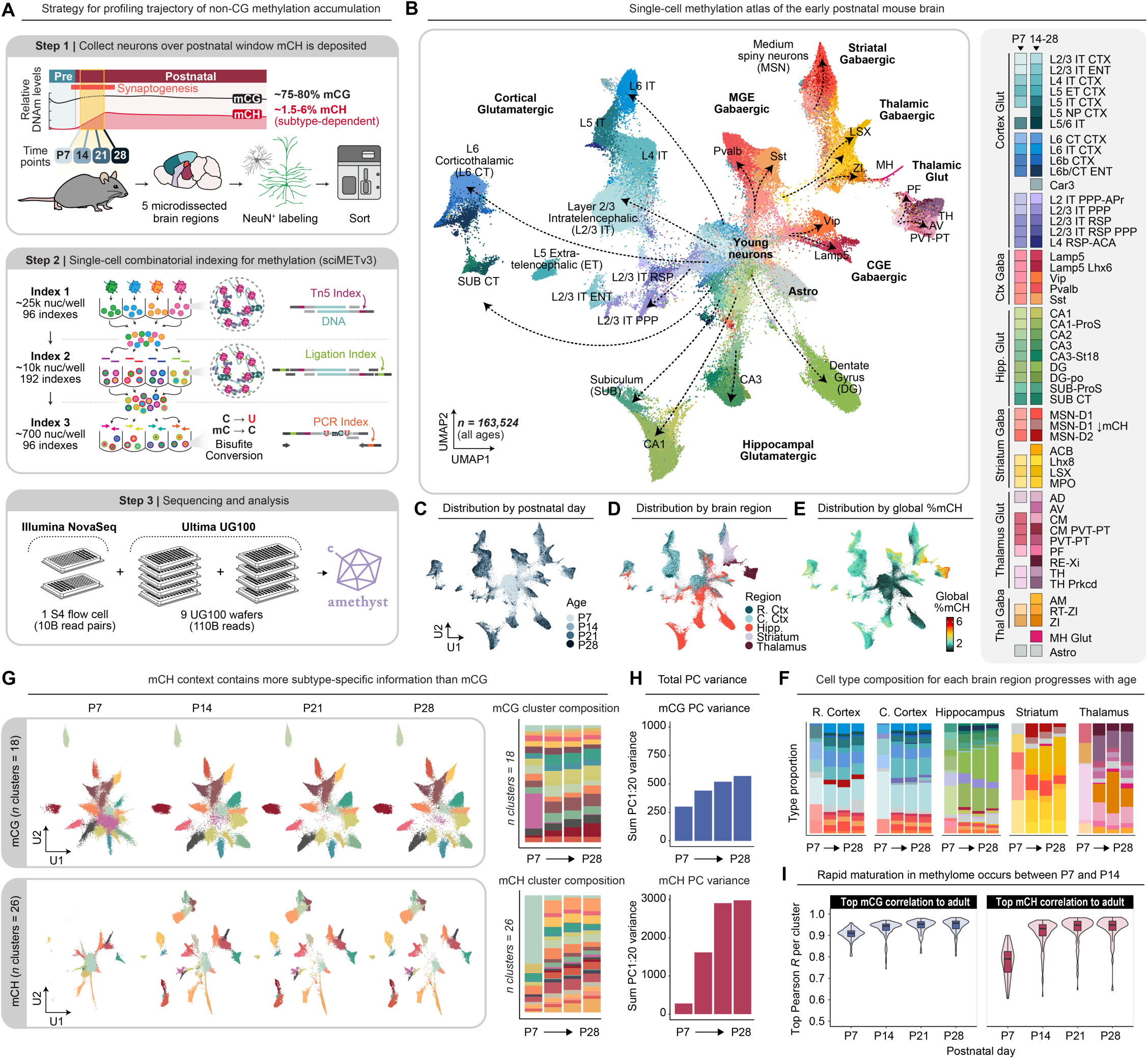
A single-cell atlas of DNA methylation in the postnatal mouse brain reveals rapid maturation across subtypes between the first and second postnatal weeks. (A) Schematic illustrating the strategy for profiling the trajectory of non-CG methylation accumulation. (B) UMAP dimensionality reduction of all control (*n* = 163,524) single-cell methylation profiles analyzed, colored by cell type. Cells are also shown colored by (C) postnatal day, (D) brain region from which the sample was collected, and (E) global %mCH. (F) Bar chart colored by cell type composition derived from each age, separated by brain region. G) UMAP dimensionality reduction generated from mCG score (see Methods) over 100kb windows (top) or mCH percent over 100kb windows (bottom) divided by age. Corresponding proportions of clusters comprising each age are shown on the right. (H) Bar chart showing the sum of variance within the first 20 principal components (PC) for dimensionality reductions derived from mCG score over 100kb windows (top) or mCH percent over 100kb windows (bottom). (I) Top Pearson *R* correlation of pseudobulked mCG (left) and mCH (right) percentages over 100kb windows between clusters in this study and adult subtype profiles.(*33, 34*)

To resolve distinct populations from billions of base-level calls, we performed dimensionality reduction using mCG and mCH levels over 100kb genomic windows (see **Methods**), a standard approach(*33–35*) that captures large-scale fluctuations in epigenomic topology while maintaining resilience to noise. Cells segregate by biological variables such as age (**Fig. 1C**), brain region (**Fig. 1D**), and global mCH (**Fig. 1E**) with minimal structure from technical features such as coverage (local kNN consistency scores of 0.63, 0.44, 0.83, 0.25, respectively). Annotation was then carefully performed for each age and subregion by consensus analysis of multiple approaches, including: cross-referencing to adult atlases,(*33*, *34*) anticorrelation of gene body mCH to RNA databases,(*95*) and assessment of mCG patterns over canonical marker genes (see **Methods**). This strategy resolved 52 distinct subtype trajectories (**Fig. 1B**), culminating in the most comprehensive DNA methylation atlas of the early postnatal mouse brain generated to date.

We next assessed the diversity of distinct methylation profiles across both methylation modalities at each time point (**Fig. 1G-H**). In the canonical CG context, populations are least distinct at P7, with each successive age progressively diversifying (**Fig. 1G-H**). Despite this, even P7 clusters demonstrate good concordance to adult atlases in the CG context (*R* = 0.91±0.004; **Fig. 1I**), facilitating subtype annotation. In the CH context, P7 clusters are poorly correlated to adult atlases (*R* = 0.78±0.009; **Fig. 1I**) due to low global mCH levels (**Fig. 1E**). Notably, a rapid CH diversification event then occurs across all subtypes between P7 and P14 (**Fig. 1G-I**), reflected in an increase in the number of clusters identified (*n =* 22 to 26; **Fig. 1G**) cumulative principal component variance (279.4 to 1618.6; **Fig. 1H**), and correlation to adult profiles (*R =* 0.78±0.009 to 0.91±0.006 **Fig. 1I**). Developmentally, this time period coincides with key events during mouse neurodevelopment like eye opening(*86*) and the GABAergic excitatory to inhibitory switch (*87*, *88*) and similar shifts have also been noted at the transcriptomic and chromatin levels.(*93*) By P21, the noncanonical context is highly correlated to adult methylation profiles (*R* = 0.93±0.006; **Fig. 1I**), with minimal variance accumulating between the third and fourth postnatal week (2909.339 to 2984.025; **Fig. 1H**). Overall, the noncanonical CH context resolved more clusters than CG using the same parameters (*n* = 18 vs 26) and exhibited higher cumulative scaled principal component variance (CG: 570, CH: 2984 at P28), in line with previous research showing this modality contains more subtype-specific information.(*27*, *33–35*, *39*, *40*) Altogether, results highlight the first to second postnatal weeks as a crucial inflection point across all brain regions in the neuronal methylation landscape, providing subtype-level resolution to prior observations.(*27*)

### Global and gene body non-CG methylation is accumulated at a subtype-specific rate

Abundant non-CG methylation is a uniquely neuronal property with exceptional subtype-specific diversity. While the global and gene-level patterns that define distinct neuronal populations have been robustly profiled in bulk(*27*) or in a snapshot at the adult stage,(*33*, *34*) the relative subtype-specific rates of mCH accumulation have not been characterized. Our single-cell and temporal atlas facilitated resolution of both brain region and subtype-specific mCH accumulation (**Fig. 2**). While the cortex and hippocampus exhibit low levels of mCH at P7 (global mCH 1.24% ± 0.04 and 1.29% ± 0.04 respectively), thalamic populations have accumulated nearly 2% global mCH (1.86% ± 0.11), exceeding P28 levels of certain populations (e.g. dentate gyrus granule cells at 1.43% ± 0.003; **Fig. 2A**). Earlier mCH in the thalamus is supported by previous whole genome bisulfite sequencing in bulk brain tissue demonstrating the order of mCH deposition occurs in the diencephalon prior to the telencephalon(*96*), roughly paralleling the order of circuit formation (*97*) and *Mecp2* expression.(*98*) In every brain region apart from the thalamus, GABAergic populations have the most mCH at P7 and maintain the highest levels throughout development (e.g., 3.15% ± 0.20 GABAergic vs 2.53% ± 0.14 glutamatergic in the P28 hippocampus). Both cortical GABAergic groups and thalamic glutamatergic neurons need to achieve an exceptional level of diversity,(*99–101*) which could be the common feature necessitating an abundant secondary level of epigenetic regulation.

**Figure 2.**
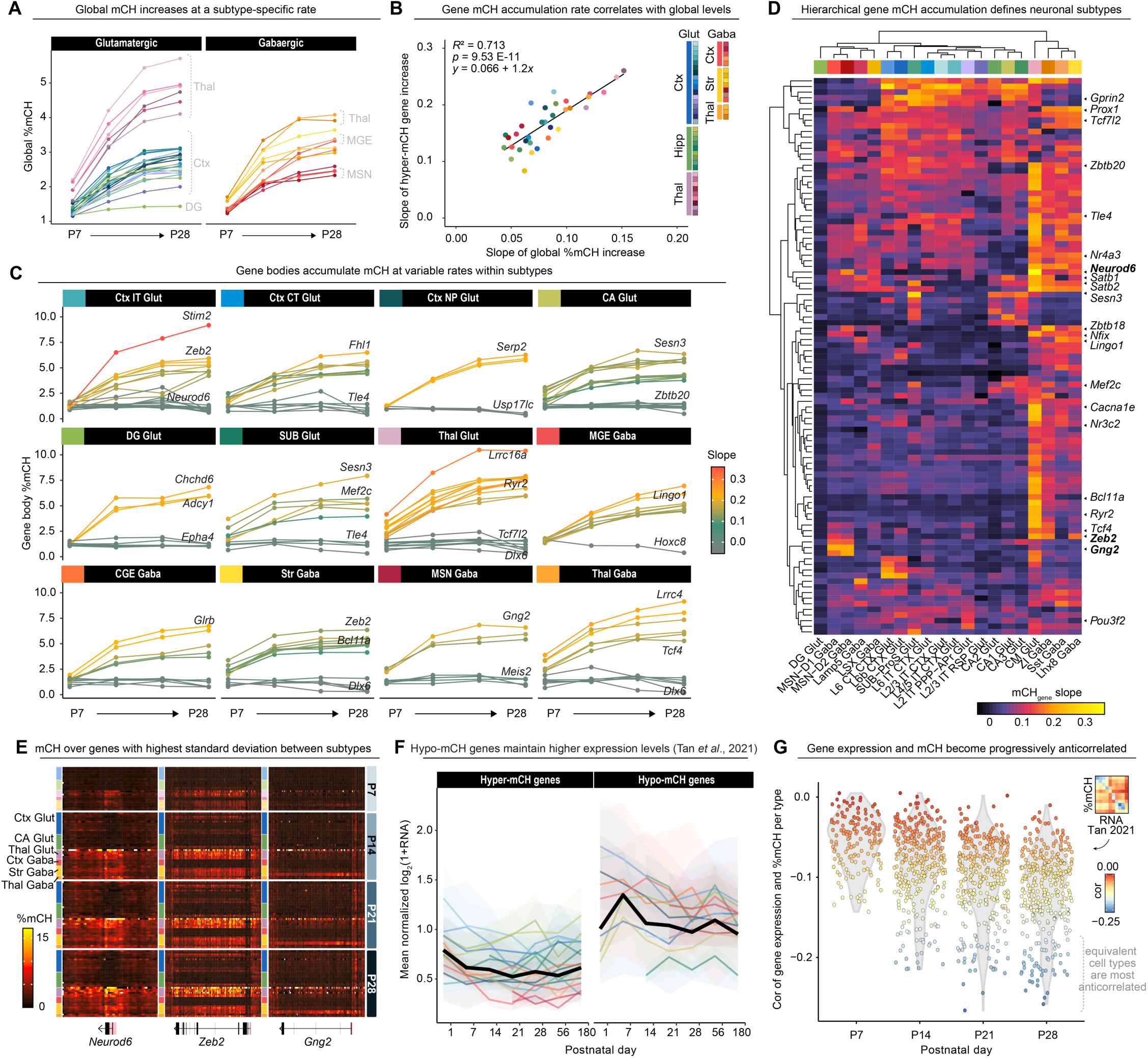
Global and gene body non-CG methylation is accumulated at a subtype-specific rate. (**A**) Global mCH values averaged by subtype at each postnatal day. Major class is annotated; subtype color corresponds to legend in Fig. 2B. (**B**) Slope of mCH accumulation on a global level (x axis) vs. for hyper-mCH genes only (y axis). (**C**) mCH accumulation trajectory for the top three genes with the highest and lowest values, faceted by major class. Lines are colored by slope. (**D**) Heatmap if subtypes hierarchically clustered by slope of mCH accumulation (color). Genes selected are the top three hypo and hyper-mCH differentially methylated genes per subtype. (**E**) Heatmap of mean mCH levels per subtype over select genes. Color represents mean mCH over 500bp windows. Gene bodies are plotted below with exons in black and putative promoter regions (start ±1500bp) in pink. (**F**) Mean normalized RNA expression from Tan *et al.*(93) of hypo and hyper-mCH genes per subtype. (**G**) Pearson R correlation of transcriptome(93) to gene mCH levels for *n =* 6,164 genes with standard deviation of >0.5 mCH between groups.

While canonical mCG displays high variability at regulatory regions, noncanonical mCH uniquely varies at boundaries of critical neuronal gene bodies, where accumulation is anticorrelated with expression.(*27*, *30*, *32–35*, *39*, *40*, *102*) Our atlas demonstrates a strong concordance between rate of mCH increase over hypermethylated genes and global mCH levels (*R*^2^ = 0.713 *p =* 9.53E-11; **Fig. 2B**). mCH accumulates steadily across targets simultaneously rather than specific genes being targeted prior to others, reflected in a maintained ratio of gene body mCH to global mCH at hypermethylated genes across timepoints (1.49±0.06). Top genes with the highest standard deviation between slopes of mCH accumulation are enriched for transcripts with critical roles in diencephalon development (*p*_elim_ *=* 2.4E-04), such as *Neurod6, Bcl11a, Satb2, Zeb2,* or *Prox1*, and genes involved in modulation of chemical synaptic transmission, such as *Gng2, Slc17a6,* or *Ryr2* (*p*_elim_ *=* 1.2E-04 **Fig. 2C-E**). Top genes with the greatest variability in mCH accumulation rates across subtypes recapitulate key candidate genes that have been profiled in detail at the adult stage,(*33*, *34*) indicating that relationships are already established by the second postnatal week.

Previous work in bulk tissue as well as parvalbumin (Pvalb) and vasopressin (Vip)-positive interneuron subtypes has demonstrated that highly expressed genes may be insulated from DNMT3A-mediated mCH accumulation via transcription-associated deposition of H3k36me3.(*27*, *30*, *32*, *103*) To assess the relationship between early postnatal gene expression and mCH levels per subtype, we leveraged a published single-cell RNA sequencing dataset of mouse cortex and hippocampus collected at identical time points.(*93*) For each subtype, we observed that hypo-mCH genes maintain higher expression levels throughout life than hyper-mCH genes (**Fig. 2F**). Overall, the variably-expressed gene RNA and mCH levels become increasingly anticorrelated from P7 onward for differentially methylated genes (*R*^2^ = 0.245; *F*(1,55) = 19.17; *p* = 5.4E-5; **Fig. 2G**). This relationship breaks down when instead comparing mCH levels to highest expressed genes (*R*^2^ = -0.017; *F*(1,55) = 0.071; *p* = 0.79), suggesting that high transcription alone does not explain low mCH regions, in agreement with previous findings in bulk tissue.(*27*) Altogether, findings support a model in which gene-level mCH is steadily accumulated on a subtype-specific basis in proportion to global mCH load, with progressive anticorrelation to gene expression.

### Synaptic partner establishment genes have a dynamic methylome in early postnatal development

To map the trajectory of CG and CH loci changes that accompany subtype maturation, we applied differentially methylated region (DMR) analysis between successive timepoint pairs. In total, we identified 366,084 CG and 19,623,286 CH DMRs across all subtypes (**Fig. 3A**). Results in both contexts again show the greatest shift between P7 and P14 with only marginal increases following, supporting findings at the 100kb level (**Fig. 1G-H**). While a similar ratio (0.99:1) of CG hyper to hypomethylations accompany subtype maturation, the CH context exclusively increases with age (112:1). Most modifications occur independently as very few loci that are hypermethylated with age have proximal changes in the reciprocal context (13.98% CG and only 0.14% CH). These observations together further highlight the distinct regulatory behaviors of the two modalities: while mCG is dynamic and can be added or removed in response to the environment, mCH is only accumulated, emphasizing its role as a stable mediator of neuronal subtype specification.(*32*)

**Figure 3.**
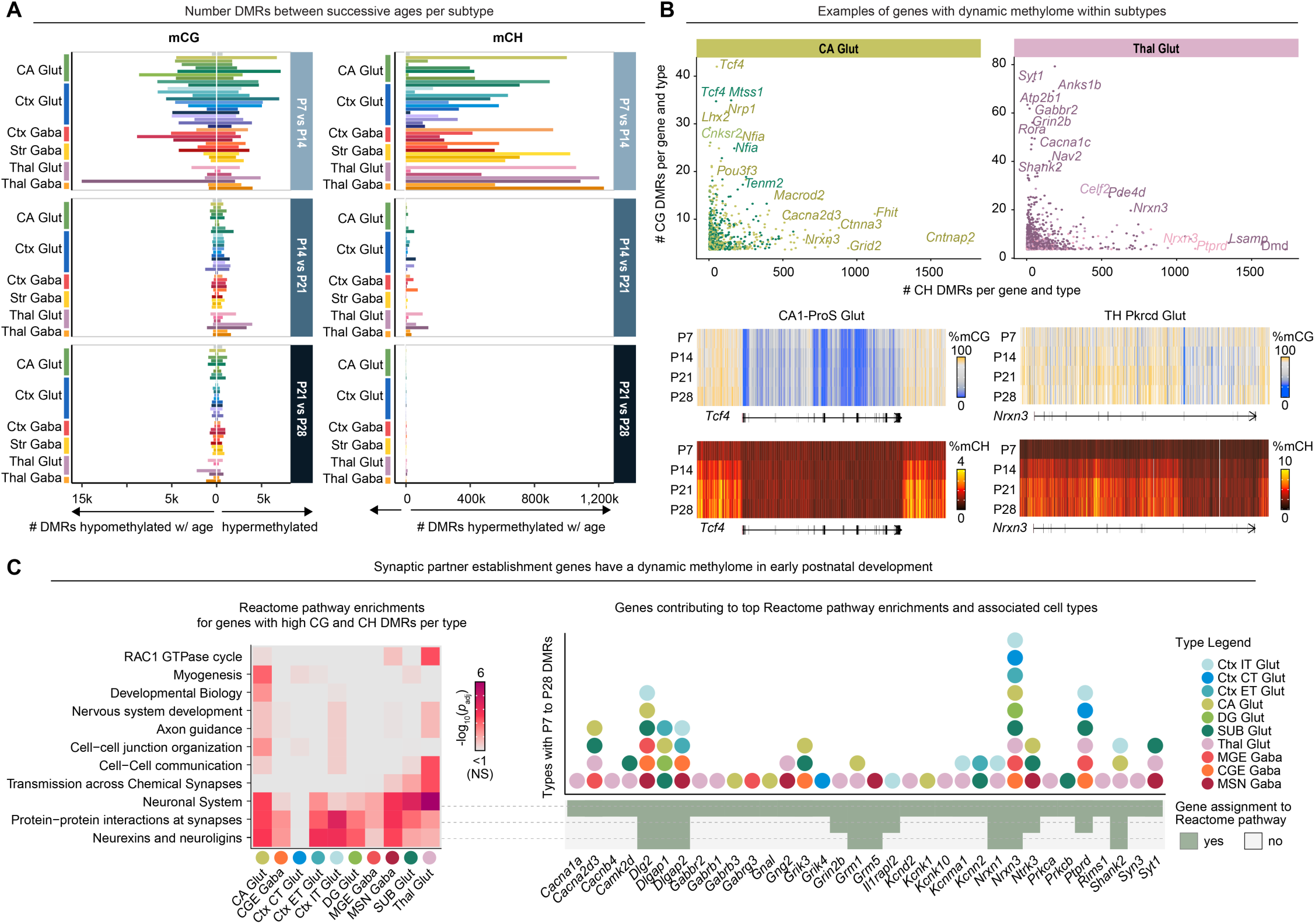
Synaptic partner establishment genes undergo dynamic methylation changes in early postnatal development. (**A**) Histogram showing number of differentially methylated 500bp windows between successive ages within each subtype. (**B**) Sum of differentially methylated 500bp windows between successive ages within subtypes for each gene. Major classes hippocampal CA glutamatergic neurons and thalamic glutamatergic neurons are used as examples. Examples of top candidates are shown in corresponding heatmaps below. Color indicates mean mCG (top) and mCH (bottom) over 500bp windows. Gene bodies are plotted below with exons in black and putative promoter regions (start ±1500bp) in pink. (**C**) Heatmap showing Reactome pathway enrichments for genes acquiring > 5 CG DMRs and > 30 CH DMRs with age. All enrichment categories identified in five or more subtypes are shown. The right panel shows all genes contributing to the top three Reactome Pathway enrichments (x axis), and in which subtypes the methylation changes with age (y axis).

We next calculated the sum of differentially methylated regions across each subtype, gene, and context to identify loci undergoing extensive methylation remodeling during development (**Fig. 3B**). Among the genes harboring the greatest numbers of DMRs were transcription factors with established roles in neuronal specification, such as *Tcf4*, *Lhx2*, and *Pou3f3* in hippocampal glutamatergic populations; or *Rora* in thalamic glutamatergic cells (**Fig. 3B**). Notably, DMR-associated genes across nearly all major classes were enriched for Reactome pathways(*104*) involving synaptic protein interactions, particularly neurexin and neuroligins (*p*_adj_ = 0.02±0.17; **Fig. 3C**), which are central organizers for both glutamatergic and GABAergic synapses in the brain.(*105*) Although the developmental onset of non-CG methylation is known to coincide with peak synaptogenesis,(*106*) the corresponding extensive methylation remodeling at genes involved in synaptic partner recognition and circuit assembly has not been observed at this resolution. For both synaptic establishment genes and neuronal transcription factors, it is paramount that proper abundances are appropriately titrated, which could evolutionarily explain the utility of an additional layer of epigenetic regulation.

### Hypomethylated region specificity associates with genomic feature composition

Regions with low methylation levels (LMRs) exhibited substantial variation in their degrees of cell type specificity (**Fig. 4A**). While distal regulatory elements are known to exhibit cell type-specific methylation patterns,(*27*, *107*) the single-cell and temporal resolution of our dataset enabled a systematic examination of the relationship between LMR specificity and underlying genomic features. We first identified all loci that exhibited low levels of methylation in at least one subtype for either methylation context (see **Methods**). The CG and CH modalities varied greatly in both the number of LMRs and proportion shared. In the CG context, we identified 156,796 loci with many regions shared across groups (28.2% in >75% of subtypes; **Fig. 4B**), in contrast to 2,202,457 CH loci with very few common between subtypes (0.56% in > 75% of subtypes; **Fig. 4C**). These observations are in line with prior understanding of mCH as containing more subtype-rich information than mCG.(*27*, *33–35*, *39*, *40*, *102*) We next examined the association of these LMRs with annotated genomic features. In the CG context, specific LMRs fall over distal enhancers (22.1%), gene bodies (35.9%), or CTCF binding sites (2.0%) - while shared CG LMRs almost exclusively fall over proximal and promoter enhancer elements (53.5% and 39.1%, respectively; **Fig. 4B**). This pattern broadly applies to the CH context but with substantially different distributions: a 2.2-fold increase in the proportion of CH LMRs was observed in intergenic, non-regulatory regions over those in the CG context (15.3% and 49.0% for CH and CG, respectively; **Fig. 5C**). This finding likely reflects megabase-scale fluctuations(*103*) in the structure in mCH patterning in addition to large regions of hypomethylation coined “mCH deserts”.(*27*) Of the CG LMRs that overlap with regulatory elements, 90.4% were also observed in CH LMRs. This suggests the two modalities are co-regulated at these hypomethylated sites and is mechanistically supported by the reduced binding affinity of DNMT3A at promoter and enhancer regions.(*32*, *103*, *108*) Furthermore, these patterns are identical across all ages, indicating conserved principles of epigenomic regulation across this dynamic reconfiguration period.

**Figure 4.**
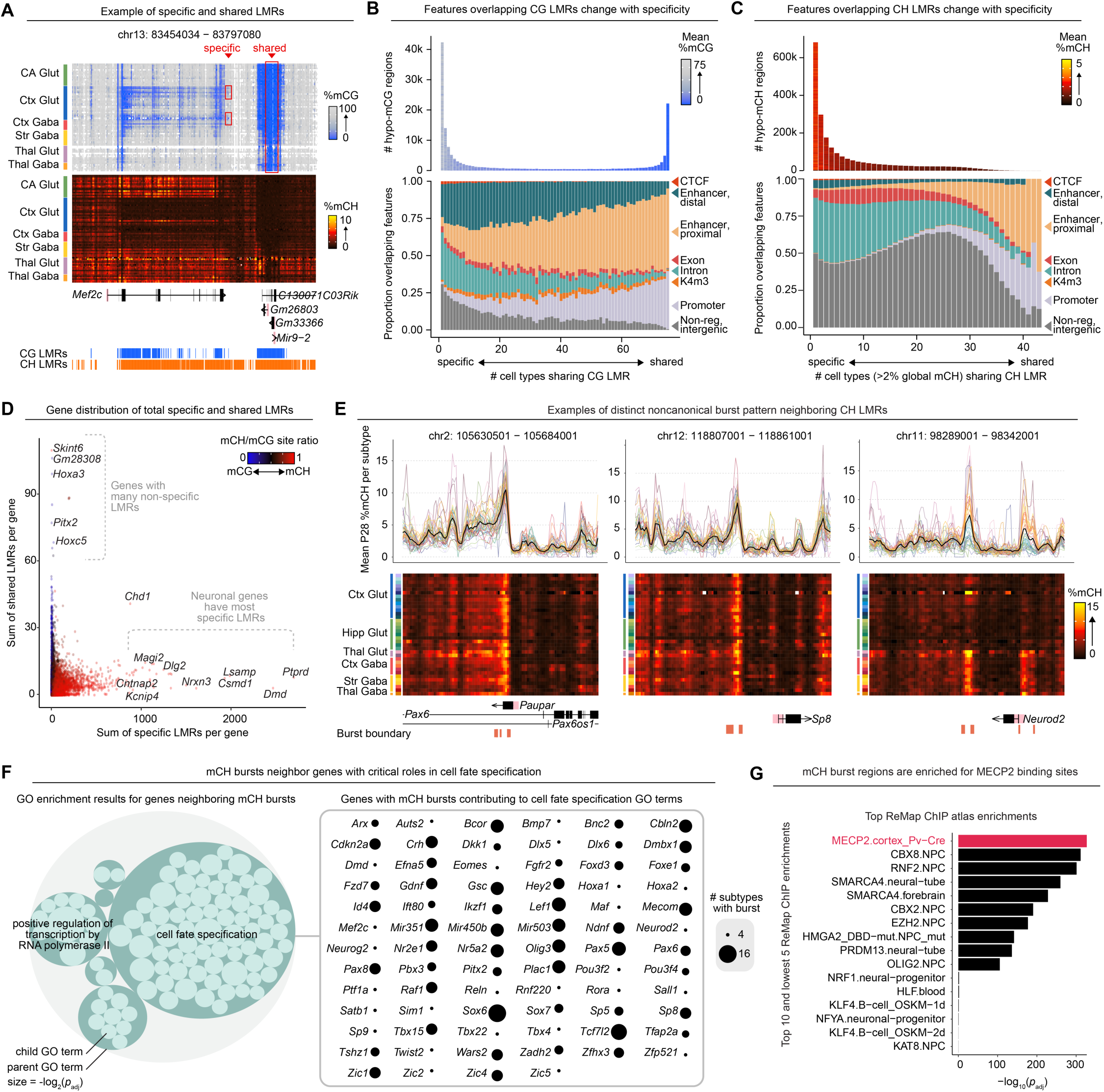
Hypomethylated region specificity associates with genomic feature composition. (**A**) Example of specific and shared LMR over *Mef2c.* Heatmaps show mean mCG (top) and mCH (bottom) per subtype over 500bp windows. Gene bodies are plotted below with exons in black and putative promoter regions (start ±1500bp) in pink. (**B-C**) Top histogram shows frequency of LMRs by subtype prevalence, where color indicates mean mCG (B) and mCH (C) across all subtypes. The bottom bar chart shows the proportion of LMRs overlapping ENCODE candidate cis-Regulatory Elements at each frequency. (**D**) Sum of specific (1–3) vs global >(*n*_groups_ - 10) LMR 500bp windows per gene. Color indicates ratio of mCH to mCG windows identified. (**E**) Examples of distinct mCH burst pattern. Mean mCH values for each 500bp window and subtype are represented as traces (top) and heatmaps (bottom). Gene bodies are plotted below with exons in black and putative promoter regions (start ±1500bp) in pink, along with the identified burst boundaries. (**F**) Circlepack diagram of all significant (*p_elim_* < 0.05) gene ontology (GO) enrichments for genes ±15000bp of mCH bursts. The circle size corresponds to -log_2_(*p_elim_*); related terms are nested together. To the right, all genes contributing to the “cell fate specification” parent term are shown, with the size of the circle indicating the number of subtypes in which the burst was identified. (**G**) Bar chart of top 10 and bottom 5 mCH burst enrichments with the ReMap ChIP atlas. Only brain and blood (control) samples were tested.

**Figure 5.**
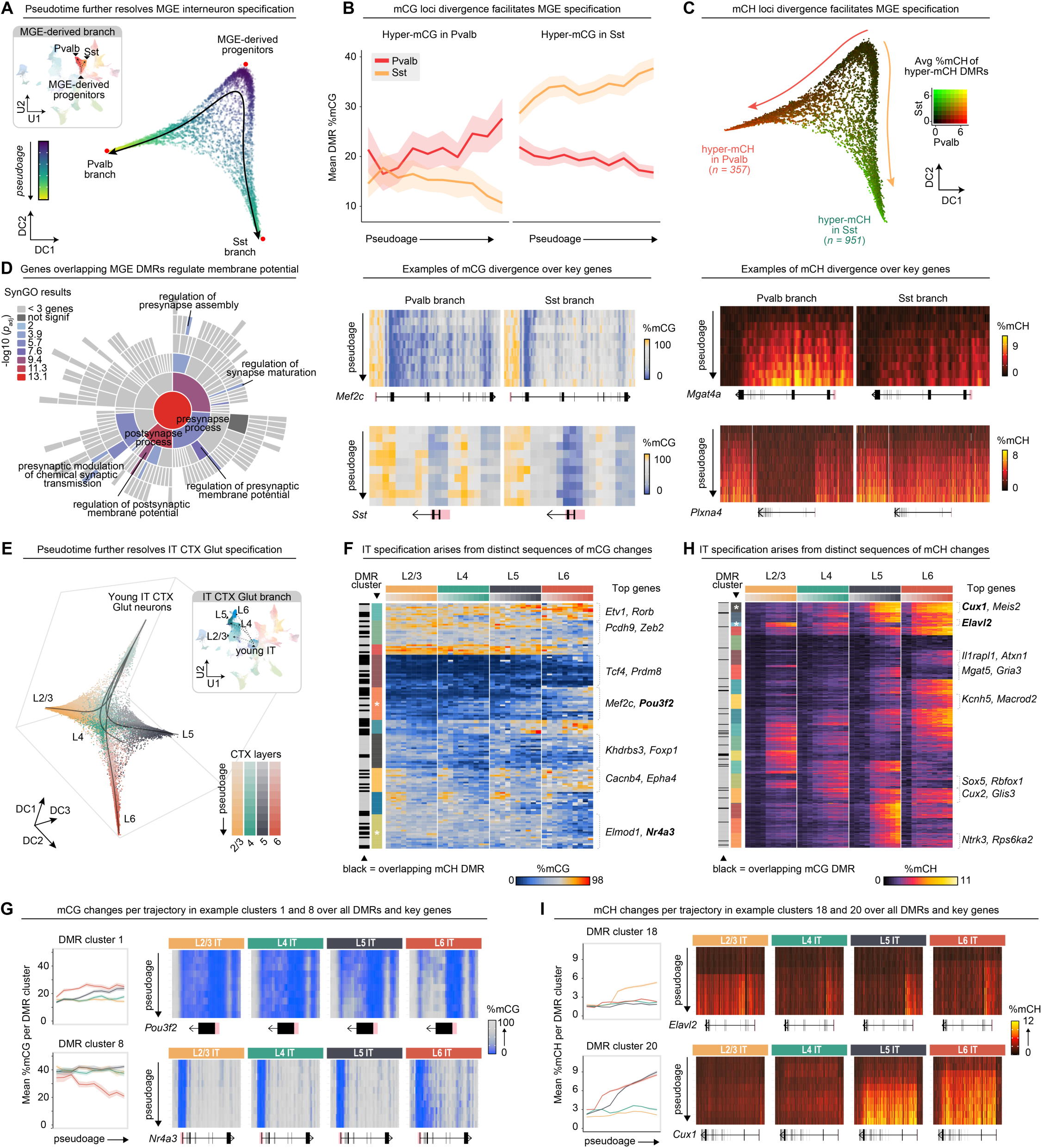
Coordinated methylation changes occur at synaptic genes regulating membrane potential during neuronal subtype diversification. (**A**) Medial ganglionic eminence-derived GABAergic neurons (MGE Gaba; highlighted in the inset panel) arranged by diffusion components 1:2. Red dots indicate trajectory tips and the arrows show the Slingshot(116) trajectory. (**B**) Mean ± SE mCG of diverged DMR loci at each pseudoage, colored by branch. Examples of overlapping genes are shown in heatmaps below with mean mCG per subtype over 500bp windows. Gene bodies are plotted below with exons in black and putative promoter regions (start ±1500bp) in pink. (**C**) Mean mCH of diverged DMR loci plotted over diffusion components. Color scale is additive with Sst hyper-mCH loci in green and Pvalb hyper-mCH loci in red. Examples of overlapping genes are shown in heatmaps below with mean mCG per subtype over 500bp windows. Gene bodies are plotted below with exons in black and putative promoter regions (start ±1500bp) in pink (**D**) Sunburst plot showing hierarchical SynGO biological process enrichment results for genes overlapping DMRs between the Pvalb and Sst branches. (**E**) Intratelencephalic cortex glutamatergic neurons (IT CTX Glut, highlighted in the inset panel) arranged by diffusion components 1:3 and colored by branch. Arrows show Slingshot trajectories. (**F**) Heatmap of hierarchically clustered DMR loci (up to top 25 per cluster). Color indicates %mCG. Pseudoage bin is on the x axis. Example genes with the most DMR overlaps are listed to the right. Clusters with an asterisk (1 and 8) are shown in more detail in (**G**) Mean mCG of each branch for DMRs in clusters 1 and 8. Shown to the right are heatmap examples of key genes in each cluster with mean mCG per pseudoage bin over 500bp windows. Gene bodies are plotted below with exons in black and putative promoter regions (start ±1500bp) in pink. (**H**) Heatmap of hierarchically clustered DMR loci (up to top 25 per cluster). Color indicates mCH. Pseudoage bin is on the x axis. Example genes with the most DMR overlaps are listed to the right. Clusters with an asterisk (18 and 20) are shown in more detail in (**I**) Mean mCH of each branch for DMRs in clusters 18 and 20. Shown to the right are heatmap examples of key genes in each cluster with mean mCH per pseudoage bin over 500bp windows. Gene bodies are plotted below with exons in black and putative promoter regions (start ±1500bp) in pink.

We next identified which genes had the most shared versus specific LMRs. Genes with the most shared LMRs include candidates not expected to vary across neuronal subtypes, such as members of the *Hox* family or T cell regulator *Skint6* (**Fig. 4D**). In contrast, genes enriched for subtype-specific LMRs play central roles in neurodevelopment and synapse assembly (GO (*p_elim_ =* 4.4E-11), including *Ptprd, Nrxn3,* and *Cntnap2*. Together, these findings support previous work(*27*, *32*, *34*, *39*) while further refining the relationship between methylation architecture across a gene and putative biological relevance. In both contexts, subtype-specific LMRs are preferentially localized to distal regulatory elements and gene bodies, whereas shared LMRs almost exclusively fall over proximal regulatory elements and promoters. This finding suggests that methylation status at proximal regulatory elements is a poor predictor of cell type-specific transcriptional output and therefore unreliable for RNA imputation.

We observed that some CH LMRs were flanked by a focal hypermethylated bursts around genes such as *Pax6, Sp8,* and *Neurod2* (**Fig. 4E**). Bursts were unique to the mCH context and did not extend to mCG. To further explore this phenomenon, we performed a genome-wide search for these bursts (see **Methods**) across each major neuronal class. 2,434 unique sites were identified in total with the majority (70.1%) neighboring a gene. 68% of GO enrichments for these genes fell under the parent category of cell fate specification due to hits such as *Pax, Sox, Sp,* and *Zic* families (**Fig. 4F**). To better understand putative functional relevance of these bursts, we tested for enrichment against all brain ChIP transcription factor binding site datasets assembled in the ReMap atlas.(*109*) Notably, the strongest enrichment (*p*_adj_ = 0) was for MECP2 binding sites in Pvalb interneurons.(*32*) This enrichment suggests a biological relevance as MECP2 is the only known mCH reader.(*53*) Collectively, these findings highlight mCH bursts as a putative mechanism through which MECP2 regulates regional specification genes following late embryogenesis, facilitating progression into the next phase of early postnatal neurodevelopment.(*110*, *111*)

### Coordinated methylation changes occur at synaptic genes regulating membrane potential during neuronal subtype diversification

Though most neuronal populations are established by P7 - albeit in an immature state - Pvalb interneurons are uniquely delayed in their specification until around the second postnatal week.(*112*) We leveraged our P7-P28 atlas to capture the methylation changes facilitating the divergence of medial ganglionic eminence (MGE)-derived interneurons into distinct Pvalb and somatostatin (Sst)-expressing populations (**Fig. 5A-D**). Principles from pseudotime analysis were applied to capture progressively resolved cell states, which strongly correlated with known age (*R* = 0.87, *p <* 2.2E-16). To identify diverging loci between the two branches, we performed DMR analysis between populations with the highest pseudoage in each lineage, identifying 281 CG sites and 1,308 CH sites in total (**Fig. 5B-C**; see **Methods**). Genes neighboring mCG DMRs include those with known roles in Pvalb interneuron specification like *Mef2c* and *Sst* itself (**Fig. 5B**).(*100*, *113*) Top DMRs for the CH context include numerous signaling receptors such as *Plxna4, Srrm4,* and *Errb4* (**Fig. 5C**), in accordance with the enrichment of synaptic partner establishment genes found previously (**Fig. 3C**). Variable methylation in such genes could prime these two classes for differential intracellular responses arising from environmental guidance cues as the interneurons migrate through the cortex to reach their final destinations. Notably, ontology terms from the SynGO database were strongly enriched for genes with roles in regulating both presynaptic and postsynaptic membrane potential (*p*_adj_ = 2.60E-06 and 4.28E-07, respectively; **Fig. 5D**), which may facilitate the fast vs. slow-spiking features that characterize these two subtypes.(*114*)

We next extended the analysis to intratelencephalic glutamatergic populations of the cortex (IT Glut CTX). Trajectory construction yielded four branches, each extending towards neurons derived from a major cortical layer (L2/3, L4, L5, L6; **Fig. 5E**). DMR analysis between the cells of oldest pseudoage yielded 876 CG and 2,476 CH loci. To better understand the relationship of changes facilitating specification of each branch, we performed hierarchical clustering (**Fig. 5F-I**), revealing both shared and distinct features of each lineage. For example, genes such as superficial layer transcription factor *Pou3f2* become progressively hypermethylated in both L5 and L6 IT Glut CTX neurons in CG DMR cluster 1, while early neuron transcription factor *Nr4a3* in cluster 8 is specifically hypomethylated only in L6 IT Glut CTX neurons. Neighboring layers are most correlated to each other with earliest born L6 having the most distinct profile. This could reflect the unique roles of L6 IT neurons, which regulate gain for both superficial circuits and corticothalamic circuits.(*115*) The contrast between branches was more striking in the CH context (**Fig. 5H**) where hierarchical clustering again resolved both shared and distinct specification patterns defining each trajectory (**Fig. 5H-I**). For example, *Elavl2* and cluster 18 DMRs are uniquely hypermethylated in L2/3 neurons, while layer 2-4 transcription factor *Cux1* is hypermethylated in L5 and L6 neurons. Though different genes neighbor IT Glut DMRs than MGE Gaba DMRs, the biological class is similar with Reactome enrichments such as regulation of postsynaptic membrane potential (*p*_adj_ = 3.54E-04) and integral component of the postsynaptic density membrane (*p*_adj_ = 2.72E-07). Altogether, these findings show that in early postnatal development, neuronal subtype diversification coincides with distinct, branch-specific methylation changes to genes that establish diverse electrophysiological properties.

### Loss of MECP2 results in an increasing number of differentially methylated regions with age and subtype-specific deficits in global mCH accumulation

The only known reader of noncanonical mCH is MECP2, loss of which causes the devastating neurodevelopmental disorder Rett syndrome. In addition to binding mCH, MECP2 has also been shown interact with DNMT3A,(*80*) facilitate topologically associated domain formation,(*36*) and organize chromatin state(*54*, *77*, *78*, *117*), all of which inform mCH deposition.(*27*, *67*, *70*) We next performed parallel analysis with mice containing a MECP2 p.W104X patient truncating variant(*89*, *90*) to interrogate how noncanonical methylation accumulation proceeds in the absence of feedback, and to explore the methylation landscape of Rett syndrome. In total, 171,845 single-cell methylomes were profiled from ten MECP2^+/y^ (*n =* 163,524) and ten MECP2^W104X/y^ (*n =* 73,714) mice spanning four ages (P7, P14, P21, P28) and two brain regions (caudal cortex and hippocampus; **Fig. 6A**; **Table S1**).

**Figure 6.**
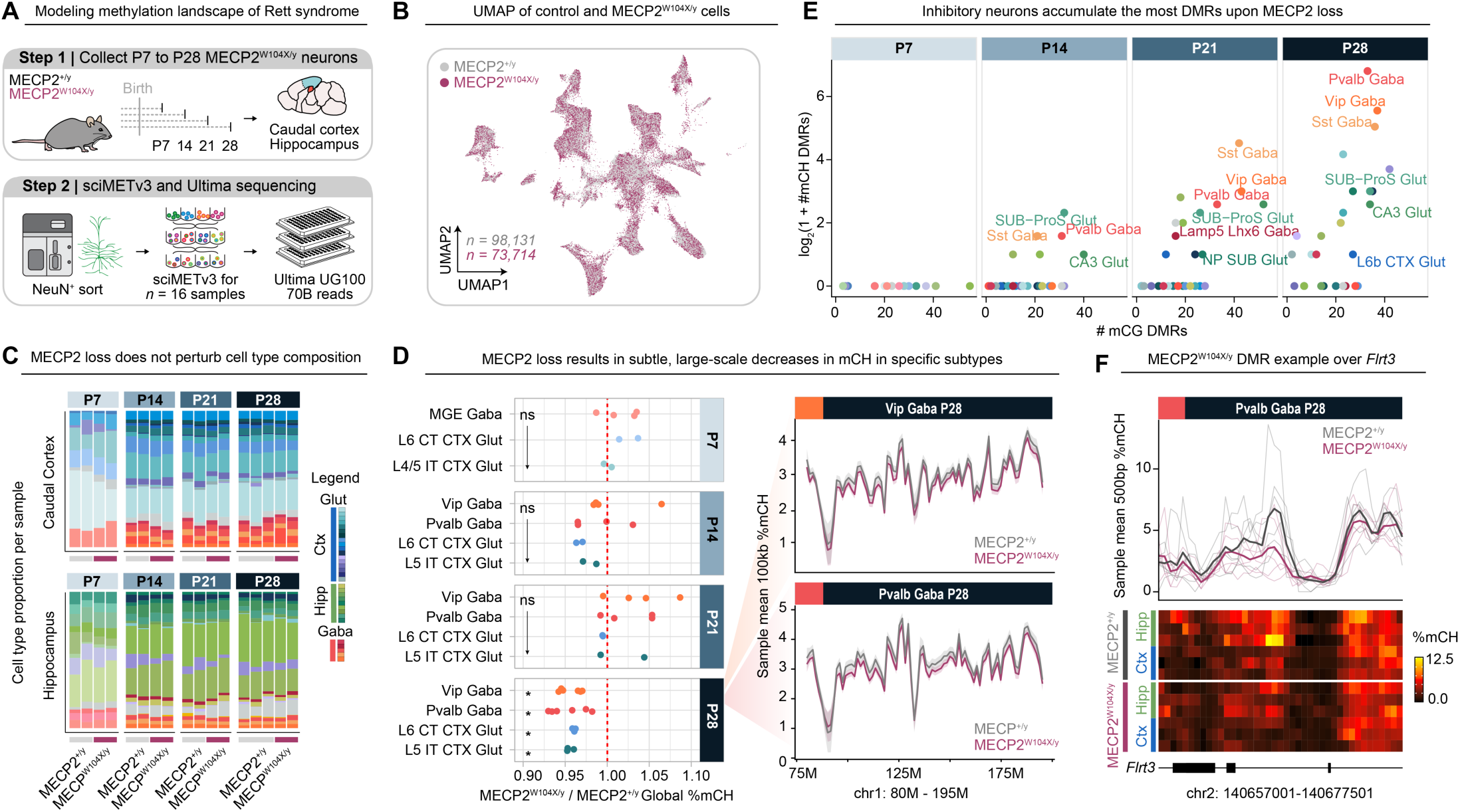
Loss of MECP2 results in an increasing number of differentially methylated regions with age and subtype-specific deficits in global mCH accumulation. (**A**) Schematic illustrating the strategy for profiling the trajectory of non-CG methylation accumulation upon MECP2 loss. (**B**) UMAP dimensionality reduction colored by genotype with *n =* 98,131 MECP2^+/y^ and *n =* 73,714 MECP2^W104X/y^ cells. (**C**) Bar chart colored by cell type composition in each age and brain region for each biological replicate. (**D**) Ratio of mean global mCH levels in MECP2^W104X/y^ normalized to MECP2^+/y^ cells per biological replicate. * *p*_adj_ < 0.05 using a two-sided *t* test with Benjamini Hochberg correction. To the right, mean mCH per biological replicate (*n =* 6 MECP2^+/y^ ; 6 MECP2^W104X/y^) over 100kb windows for chr1:80000000-195000000 in P28 Vip Gaba and Pvalb Gaba cells. (**E**) Total consensus DMRs (see **Methods**) in MECP2^W104X/y^ cells for each age and subtype. The y axis is on a log_2_(1 + x) scale. (**F**) Example of CH DMR in P28 Pvalb Gaba cells over the *Flrt3* gene. Top trace shows mean %mCH per biological replicate (*n =* 6 MECP2^+/y^ ; 6 MECP2^W104X/y^) over 500bp windows. The bottom panel shows this information as a heatmap with one row per replicate. *Flrt3* is mapped below with exon location indicated by the black rectangles.

We first assessed whether any large-scale shifts in the methylome could be identified upon loss of MECP2. No distinction was observable in dimensionality reductions between MECP2^+/y^ and MECP2^W104X/y^ cells (silhouette = 0.003; **Fig. 6B**). Though single-cell analyses have limited utility as cell counters, subtype proportions are similar between genotypes (**Fig. 6C**), in line with previous findings using *in vitro* models.(*75*) Global CG levels are not different at any age or subtype. Onset of CH accumulation also is not perturbed upon MECP2 loss, with no difference detected in global mCH levels in any subtype at age P7 to P21. However, four classes of neurons failed to reach control global mCH levels in P28 MECP2^W104X/y^ cells (**Fig. 6D**): Vip Gaba (*p*_adj_ *=* 0.014), L6 CT CTX Glut (*p*_adj_ *=* 0.014), L5 IT CTX Glut (*p*_adj_ *=* 0.032), and Pvalb Gaba (*p*_adj_ *=* 0.032). This reduction presents as a consistent shift across extensive regions of the genome in all four populations (**Fig. 6D**). While modest (4.36%±0.16%), this corresponds to a change in methylation status of ∼ one million CH sites. Such a widespread effect is in line with the subtle, global phenotypes that are typically described upon MECP2 loss, due its ubiquitous binding across the genome.(*54*, *82*)

Focal DMR analysis identified 2,472 genotype-associated DMRs across cell types. Consistent with the global reduction in mCH observed in **Fig. 6D**, most DMRs (70.8%) were hypomethylated in MECP2^W104X/y^ cells relative to controls. The number of DMRs increased progressively with age following MECP2 loss, particularly in GABAergic interneurons (**Fig. 6E**). Notably, this trajectory parallels the developmental accumulation of both MECP2 and non-CG methylation, which reach higher levels in cortical GABAergic populations relative to glutamatergic.(*33*, *40*, *59*) Among all subtypes, Pvalb interneurons have the most DMRs upon MECP2 loss, consistent with extensive evidence implicating this population as a key contributor to Rett pathology.(*30*, *59*, *81*, *118–122*) Genes associated with DMRs in Pvalb MECP2^W104X/y^ cells are enriched for processes relating to axonogenesis (*p_elim_* = 8.5E-04; 34.3% of GO terms), regulation of neuron projection development (*p_elim_* = 6.5E-03; 14.9%); and cellular divalent inorganic cation homeostasis (*p_elim_* = 8.0E-03; 6.3%). These enrichments were driven by DMRs such as chr2:140667001-140667501 (*p*_adj_ *=* 0.013) over *Flrt3* (**Fig. 6F**), which encodes a cell adhesion molecule that mediates the tangential migratory stream of cortical interneurons during development.(*123*) Across DMR GO enrichments for all cell types, neuron projection development (*p_elim_* = 0.016±0.004; 15.7% of subtypes) and synaptic signaling (*p_elim_* = 0.03±0.003; 13.1% of subtypes) occurred most frequently. Together, these findings suggest that MECP2 loss preferentially disrupts methylation at genes involved in neuronal connectivity and circuit formation. Given the central role of these pathways in neurodevelopment, even modest perturbations to their regulatory landscapes may contribute to the widespread dysfunction that is characteristic of Rett syndrome.

## Discussion

In this study, we have generated the most comprehensive single-cell DNA methylation atlas of the early postnatal mouse brain to date. This resource enabled reconstruction of both canonical (CG) and noncanonical (CH) methylation dynamics underlying the specification of 52 distinct neuronal subtypes, providing temporal context to single-cell methylation atlases at the adult stage.(*33–35*, *40*, *124*) Our analyses consistently highlight between the first and second postnatal week as a critical inflection point over which methylome profiles rapidly mature towards an adult-like state. Findings provide further resolution to what has been shown in bulk tissue,(*27*, *38*) showing that this transition occurs across brain regions and subtypes, most prominently in the noncanonical context at genes involved in synaptic partner establishment. Comparable transitions during this window have also been reported at the transcriptomic and chromatin levels,(*93*) implicating an epigenome-wide remodeling event. Developmentally, this window coincides with many defining milestones - such as eye opening,(*86*) peak synaptogenesis onset,(*125*) the GABAergic excitatory to inhibitory switch,(*87*, *88*) and the GABA-driven transition from synchronized to sparse neural activity.(*126–128*) These observations collectively highlight the first to second postnatal week as an extremely dynamic period in the life of a postmitotic neuron while it integrates into functional circuits. Since proper circuit formation is both extraordinarily complex and highly sensitive to perturbation - as evidenced by autism, epilepsy, and a host of other neurodevelopmental disorders, our observations support a model in which non-canonical methylation provides an additional layer of epigenetic regulatory control over crucial neuronal genes involved in maturation and connectivity.(*30*, *36*, *103*) Given the prominence of Eph/ephrin, Ntrk, and Nrxn signaling receptor families among the most dynamically methylated genes, an intriguing future direction would be to assess the relationship of autonomous methylation changes and those initiated by guidance cues, transcription factor gradients, and other non-autonomous signals uniquely found in the developing brain.

The clear relationship between non-CG methylation and Rett syndrome has prompted multiple other investigations of the methylation consequences upon MECP2 loss using either bulk striatal tissue(*67*) or purified populations(*30*, *36*, *129*) in conjunction with bulk-cell profiling. In line with Rett syndrome literature, results have been mixed and subtle.(*82*) Our single-cell approach harmonizes some of these apparent discrepancies and paints a more complete picture of the methylation landscape of Rett syndrome. For example, findings corroborate subtype-specific, smaller focal DMRs enriched in interneurons, in agreement with Jin and colleagues’ report of more DMRs in pooled GABAergic versus glutamatergic populations.(*129*) Our finding of increased DMRs in GABAergic neurons with age is notable given that selective deletion of MECP2 in these populations has been shown to recapitulate most Rett symptoms,(*59*, *81*) including motor deficits, which are pronounced in this model.(*90*) Results also support no global methylation differences found upon MECP2 loss in many subtypes - such as medium spiny neurons.(*67*) However, we do observe subtle, yet significant, global mCH decreases in four populations, including parvalbumin-positive interneurons, highlighting the power of the single-cell approach. A putative mechanism for this intriguing shift in mCH but not mCG could be explained by the relationship between synaptic activity and noncanonical methylation accumulation. An abundance of evidence points towards a positive association between synaptic activity and mCH deposition(*27*, *32*, *96*, *130*, *131*) and mice lacking MECP2 have reduced synaptic strength and density.(*132*) Future studies could use a *Mecp2* duplication model to investigate this relationship further.

Rett syndrome is a compelling candidate for future study because of its potential for effective therapeutic intervention. Though it is a regressive disorder accompanied by both neuronal and glial dysfunction, many molecular and functional phenotypes are importantly ameliorated upon re-introduction of MECP2 in either neurons or glia.(*90*, *133–136*) However, achieving expression levels that closely approximate endogenous MECP2 is critical for successful rescue,(*46*) which is why variants amenable to RNA editing, such as W104X in the mouse model, hold valuable therapeutic potential.(*46*) An important future direction for RNA editing would therefore be to determine whether ADAR2-mediated correction of the W104X truncation, which has been shown to rescue respiratory phenotypes, recovers mCH levels in parallel.(*90–92*, *137*) Given the developmental timeline of mCH accumulation, the efficacy of MECP2 reintroduction for correcting methylation phenotypes may have a parallel critical window. Future studies should assess if there is persisting mCH loss when MECP2 is re-introduced in adulthood and whether that contributes to residual functional deficits.(*135*) Furthermore, while experiments in this study were performed in MECP2-null male mice to better understand the role of non-CG methylation in neurodevelopment, an important next step is to extend this analysis to females, which are more analogous model to the human disorder.(*44*, *47*) Properties of random X-inactivation simultaneously provide both the advantage of intra-individual controls and the bioinformatic challenge of identifying MECP2-deficient cells. While standard library preparation approaches do not facilitate sufficient coverage over the variant-containing region to determine X-inactivation status, a targeting strategy could readily be implemented to help deconvolute which allele is expressed.(*138*)

In addition to raising many intriguing avenues for future direction, this work has laid the foundation for defining the early postnatal methylation dynamics that facilitate neuronal subtype specification. Our single-cell strategy provides unprecedented insight into this process as well as the epigenetic basis of Rett syndrome, collectively implicating methylation as an underlying contributor of the ubiquitous transcriptomic dysregulation that has made Rett pathology so challenging to understand. As single-cell methylation technologies become more widely accessible, previously intractable questions can be addressed, enabling a deeper understanding of the epigenetic landscape of mammalian neurodevelopment across both healthy and diseased contexts.

## Materials and Methods

### Generation of *Mecp2^G311A^*^/+^ knock-in mouse strain

The mouse lines used in this study were first described by Sinnamon and colleagues(*90*) as follows. The database https://chopchop.cbu.uib.no/was used to scan *Mecp2* to identify sites to generate the *Mecp2* 311G>A; MECP2 W104X knockin mutation. The *Mecp2* sgRNA (5’- ACC CCA CCT TGC CTG AAG GT) was prepared by *in vitro* transcription using MEGAshortscript T7 kit (Life Technologies) and a gBlock fragment from IDT as template. The gBlock fragment contains the T7 promoter followed by MECP2 sgRNA sequence, crRNA, connecting loop and tracRNA. Transcribed sgRNA was then purified using a MEGAclear kit (Life Technologies). To prepare sgRNA/Cas9 ribonucleoprotein, sgRNA was incubated with Cas9 protein (obtained from IDT) at a final concentration of 50 ng/ul and 150 ng/ul, respectively. A single-strand DNA donor (obtained from IDT), which was used as template for homology-directed repair to introduce the *Mecp2* 311G>A; MECP2^W104X^ mutation, was mixed with the sgRNA/Cas9 ribonucleoprotein to the final concentration of 50 ng/ul. The donor sequence is: 5’- A*A*G* ACT TGC TCT TAC TTA CTT GAT CAA ATA TAC ATC ATA CTT TCC AGC AGA TCG GCC AGA CTT CCT TTG TTT AAG CTT TCG TGT CTA ACC TTC AGG CAA GGT GGG GTC ATC ATA CAT AGG TCC CCG GTC ACG GAT AAT GGA GCG CCG* C*T*G (T is the 311G>A knockin mutation and *denotes phosphorothioate linkages to prevent degradation). This mixture was then microinjected into the pronucleus of C57BL6 (Jackson Laboratory #000664) one-cell embryos and then transferred into pseudo pregnant females for subsequent development. Founder animals were initially identified by PCR using primer pair (nMC-W104XF: 5’- CCC ACC TTG CCT GAA GGG TA and nMC-I3R: 5’- CCT AGC CTT CCT ACC CAC CTG) that amplified a fragment of 146 bp specific to the *Mecp2* G311A; MECP2 W104X knock-in mutation. Potential founders were further confirmed by PCR using primer pair (nMeCE3F: 5’- CTC AGG CTC TGC CCC AGC AG and nMC-I3R: 5’- CCT AGC CTT CCT ACC CAC CTG) to amplify a fragment of 237 bp spanning the knock-in mutation followed by sequencing of the PCR product to confirm their identity.

### Animal care and tissue acquisition

All animal procedures were approved by the Institutional Animal Care and Use Committees of Oregon Health and Science University under IRB protocol #TR02_IP00000284. All mice were housed under specific-pathogen-free conditions and controlled humidity (target value 55%, minimum 45%, maximum 65%), temperature (target value 22 °C, minimum 20 °C, maximum 24 °C) and lighting (12/12-h light/dark period) and free access to food (regular mouse chow) and water. The *Mecp2^G311A/+^* mice are maintained by crossing to pure wild-type C57BL/6J mice (Jax #000664). Genotyping was performed using primers specific for the *Mecp2^G311A^* allele (Fwd: CCCACCTTGCCTGAAGGGTA; Rev: CCTAGCCTTCCTACCCACCTG). Separately, sex was determined using PCR primers specific for the X and Y chromosomes (Fwd: CACCTTAAGAACAAGCCAATACA; Rev: GGCTTGTCCTGAAAACATTTGG). Brain tissue was collected at four postnatal ages: P7, P14, P21, and P28. Mice were euthanized by brief exposure to isoflurane before decapitation. All brain subregions were dissected according to boundaries described in the Allen Brain Atlas database. Tissue samples were placed in ice cold HBSS with 10mM HEPES solution. Subregions were then snap frozen using liquid nitrogen before storage at -80°C until needed.

### Isolation of neurons from dissected mouse brain tissue

To isolate neurons from dissected mouse brain tissue, frozen samples were first thawed in a solution of nuclei isolation buffer containing Triton X-100 (TXNIB), made as follows: 0.32M sucrose, 5mM CaCl_2_, 3mM MgOAc (Sigma Aldrich #M5661), 10mM HEPES-KOH pH7.2 (Fisher Scientific #BP 310-1), 0.1% TX-100, 1X protease inhibitors (Fisher Scientific #PIA32955). Samples were triturated using a 1 mL pipet and passed through a 40 μM Flow-Mi strainer (Millipore Sigma #136800040) to generate a single-nucleus cell suspension. Nuclei were spun (500xg, 4°C, 5 minutes) and resuspended in 10% NGS blocking buffer (Life Technologies #500622) and 0.1% TX-100. Nuclei were blocked for 15 minutes at 4°C on a shaker in the blocking solution. 1:500 anti-NeuN antibody conjugated to AlexaFluor 488 (Sigma Aldrich #MAB377X:488) was then added to the cell suspension in blocking buffer. Nuclei were further incubated shaking at 4°C for 1.5 hours. DNA was stained using a brief (∼10 minute) incubation with DAPI (5 μg/mL; Fisher Scientific #D1306). After staining, up to 1 million DAPI+ NeuN+ nuclei per subregion were selected using a Sony SH800 flow sorting instrument. Collection tubes contained 100 μL NIB (10 mM HEPES, 10 mM NaCl, 3 mM MgCl_2_, 0.10% Igepal, 0.10% Tween-20) to help prevent bursting of the fragile nuclei during sorting. Isolated neurons were spun (500xg, 4°C, 5 minutes) and fixed in 476 μL NIB + 23.5 μL 16% formaldehyde (Fisher Scientific #PI28906) for 10 minutes at room temperature. The solution was quenched by adding 23.5 μL of 2.5M glycine (Sigma-Aldrich #G8898-500G) and incubated for 5 minutes further on ice. Samples were spun (500xg, 4°C, 5 minutes) and stored at - 20°C overnight in Furlong buffer (40% glycerol, 50 mM HEPES, 5 mM MgOAc, 0.1 mM EDTA, 5 mM DTT, 0.4X protease inhibitors) before carrying out the sciMETv3 protocol within 1-2 days.

### Single-cell combinatorial indexing for methylation analysis (sciMETv3) on isolated mouse neurons

#### Nucleosome disruption and tagmentation (barcode 1)

The sciMETv3 protocol(*83–85*) was then applied to isolated mouse neurons. Samples were spun (500xg, 4°C, 5 minutes) out of Furlong buffer (40% glycerol, 50 mM HEPES, 5 mM MgOAc, 0.1 mM EDTA, 5 mM DTT, 0.4X protease inhibitors) and resuspended in 970 μL NIB (10 mM HEPES, 10 mM NaCl, 3 mM MgCl_2_, 0.10% Igepal, 0.10% Tween-20). Nucleosomes were disrupted by adding 30 μL 10% SDS for 20 minutes at 37°C. SDS was spun out at room temperature (5 min, 500xg). Nuclei were counted and tagmented at ∼ 20,000 nuclei / well in a 96-well plate (Fisher Scientific #AB2396) using Tn5 loaded with adapters containing all methylated cytosines (Scale Biosciences #941770). Each well contained 10 μL tagmentation buffer (Scale Biosciences #941788). The plate was incubated at 55°C for 15 min and then placed on ice. Following tagmentation, nuclei were pooled and washed twice: first with 2 mL cold NIB, then with 3 mL cold NIB + 3 μL (New England Biolabs #B9200S). Nuclei were spun (500xg, 4°C, 5 minutes) and resuspended in 110 μL NIB, and quantified using a K2 Cellometer (Revvity #CMT-K2-MX-150).

#### Ligation (barcode 2)

To the 110 μL of nuclei, the following was added: 33 μL 10X Polynucleotide Kinase Buffer, 33 μL 10 mM ATP, 22 μL dH_2_O (Fisher Scientific #10977023) and 132 μL T4 Polynucleotide Kinase (New England Biolabs #M0201L). The solution was mixed by pipetting and distributed to a plate at 3 μL per well. The plate was incubated at 37°C for 30 min and then placed on ice. 2 μL of 15 μM ligation barcode oligos were added to each well of the plate (IDT, see Nichols *et al.,* 2024 for sequences). The following was then added to each well of the plate: 6.2 μL 2X StickTogether Buffer, 0.3 μL 100 μM v3 ligation splint and 1.5 μL T7 DNA Ligase (New Englad Biolabs #M0318L). The plate was incubated either at 25°C for 1 hr (original protocol) or 15°C overnight (later iteration) and then placed on ice and allowed to cool fully.

Post-ligation, all wells were pooled into a 5 mL tube. 3 mL NIB-H and 3 μL BSA were added. Nuclei were then spun down at 4°C 500xG for 5 min. The supernatant was removed. 3 mL NIB-H (with no protease inhibitors) was added. The tube was then spun (500xg, 4°C, 5 minutes) and resuspended in 100 μL NIB-H (no protease inhibitors). Nuclei were quantified and diluted to 375 nuclei per μL in NIB-H. Storage plates were prepped with each well containing 2 μL cell solution (750 nuclei total), 1 μL M-Digestion Buffer (Zymo Research #D5021-9), 0.07 μL Qiagen Proteinase K, and 0.93 μL dH_2_O. The plates were spun down briefly and frozen at −20°C.

#### Bisulfite conversion

Detection of methylation was performed using a bisulfite conversion-based approach. Plates were defrosted and spun down briefly to collect the liquid to the bottoms of the wells. The plates were then incubated at 50°C for 20 minutes to digest the nuclei and reverse cross-links. One bottle of CT Conversion Reagent (Zymo Research #D5003-1) was reconstituted by adding 7.9 mL M-Solubilization Buffer (Zymo Research #D5021-7) and 3 mL M-Dilution Buffer. The bottle was shaken vigorously until the solid was dissolved. Once dissolved, 1.6 mL M-Reaction Buffer (Zymo Research #D5021-8) was added and shaken vigorously. To each well of the plate of digested nuclei, 23 μL reconstituted CT Conversion Reagent was added and pipette mixed 5 times. The plate was spun down briefly. It was then incubated at 98°C for 8 minutes, followed by 64°C for 3.5 hours, with a final 4°C hold overnight.

#### Post-bisulfite conversion cleanup

Post-conversion, bisulfite reagent was cleaned. Plates were briefly spun. 120 μL M-Binding Buffer (Zymo Research #D5040-3) was added to each well of the plate and pipette mixed 5 times before transferring the entire volume to a 96-well Zymo-Spin I-96 Shallow-Well Plate (Zymo Research #C2004-SW). A custom vacuum manifold was used to pull through the M-Binding Buffer on an Agilent Bravo liquid-handling robot. Next, 400 μL M-Wash Buffer (Zymo Research #D5040-4) was added to each well and pulled through using the vacuum manifold. The vacuum was then turned off and 50 μL L-Desulphonation Buffer (Zymo Research #D5030-5) was added to each well and incubated for 15 minutes at room temperature. The buffer was then pulled through on the vacuum manifold. The plate was covered with an air permeable seal (Zymo Research #C2011-8) and spun down at maximum speed for 10 minutes to completely dry it. Elution parameters varied depending on whether the splint ligation or linear amplification method was used (see below). The protocol used is contained in the metadata for each cell.

#### Splint ligation protocol through indexing PCR (Barcode 3)

The splint ligation (SL) protocol was used for pilot analyses as it is cheaper and faster, but at a small cost in final library complexity. Buffer EB (Qiagen #19086) was preheated at 55°C. 10 μL preheated Buffer EB was added to each BSC cleanup column and the plate was re-sealed with the air permeable seal (Zymo Research #C2011-8). The column plate was put on top of an elution plate, incubated together at 55°C for 4 minutes, then spun at max speed for 10 minutes to elute.

For splint ligation, the plate was spun down and reagents - except the enzymes, 100 mM ATP, and 50% PEG 8000 - were allowed to equilibrate at room temperature. The enzymes and ATP were left on ice. The 50% PEG 8000 was heated to 55°C, and the required aliquot was placed into a new master mix tube. The SL Master Mix was assembled by adding reagents to the room temperature PEG tube. The enzymes were added last, just before pipetting to the receiving plate. The following recipe is for one 96-well plate: 180 uL 50% PEG 8000, 107.4 uL 1,3 propanediol, 82.5 uL SCR Buffer, 11 uL 1M DTT, 11 uL 100 mM ATP, 13.75 uL T4 PNK (10,000 U/mL) and 13.75 uL T4 DNA Ligase (2,000,000 U/mL; New England Biolabs #M0202M).

An ice bucket was prepared at the thermocycler, and the plate was heat-shocked at 95°C for 3 minutes and then placed on ice for 2 minutes. The plate was quickly spun down at 4°C, and 1 µL 1.5 µM pre-annealed P5 adapter (from Nichols et al. 2018) was added to each well. The plate was removed from the ice and placed at room temperature. If not already done, the PNK and Ligase were added to the SL Master Mix, which was then pulse-vortexed at max speed several times with spin downs in between. 3.8 µL room temperature SL Master Mix was added to each well using P20 tips and pipetting slowly. The plate was sealed and placed on the plate shaker at ∼1000 RPM for 5 seconds, then quickly spun down. The plate was incubated at 37°C for 45 minutes and then at 65°C for 20 minutes to inactivate the ligase. The plate was then spun down prior to indexing PCR.

The indexing PCR was performed with the following recipe for each well of a 96-well plate: 10 μL 5X VeraSeq GC Buffer, 2 μL 10 mM dNTPs (New England Biolabs #N0447L), 1.5 μL VeraSeq ULtra Polymerase (Qiagen #P7520L), 24 μL dH_2_O, 0.5 μL EvaGreen 100X (Biotium #31019) and 1 μL 1 μM i7 Flow Cell primer for a total volume of 39 μL. 1 μL of barcoded i5 primers was added separately to each well. A full list of primers can be found in Nichols *et al.,* 2024. The plate was mixed and placed on a qPCR with the following thermal conditions: 98°C initial denaturation for 30 s, 98°C for 30 s, 57°C annealing for 20 s, 72°C extension for 20 s, 72°C plate read for 10 s (these last 4 steps were cycled until exponential amplification was seen; roughly 11-13 cycles).

#### Linear amplification protocol through indexing PCR (Barcode 3)

The linear amplification version of the protocol was predominantly used as it creates higher complexity libraries, but at a higher labor and monetary cost. Buffer EB (Qiagen #19086) was preheated at 55°C. 25 μL preheated Buffer EB was added to each BSC cleanup column and the plate was re-sealed with the air permeable seal. The column plate was put on top of a pre-assembled elution plate containing 17.8 µL dH_2_O, 5 µL 10X NEB Buffer r2.1 (Fisher Scientific #B6002S), 2 µL 10 mM dNTPs (New England Biolabs #N0447L), 0.2 µL 100 µM 9H Random Primer, and 1 uL 40 ng/uL ET-SSB (New England Biolabs #M2401S) per well. The plates were incubated together at 55°C for 4 minutes, then spun at max speed for 10 minutes to elute.

The linear amp (LA) plate was then heat shocked at 95 °C for 45 s and then placed on ice until cool. 10 units of Klenow exo− (Qiagen Beverly #P7010-LC-L) was added per well and placed on a thermocycler with the following program: 4 °C for 5 min, ramp of 1 °C every 15 s until 37 °C is reached, incubation for 90 min at 37 °C, then hold at 4 °C. This is the first LA cycle. The plate was then heat shocked again at 95 °C for 45 s and placed on ice. The following was then added to each well of the plate: 0.1 µL 100 µM 9H Random Primer, 1 µL 10 mM dNTPs (New England Biolabs #N0447L), 0.5 µL 10X NEB Buffer 2.1, 1.65 µL dH2O and 2 µL of Klenow exo- (5U/μL). The above thermocycling protocol was run again. This is the second LA cycle. This cycle was repeated twice more for a total of 4 LA cycles. After completion of the last cycle, a 1.1:1 (SPRI to template ratio) SPRI cleanup was performed using Mag-Bind® TotalPure NGS beads (VWR # 75877-716) to concentrate and clean the sample. The final elution of the SPRI clean was done with 21 µL Buffer EB (Qiagen #19086) per well.

The final combinatorial barcode sequence is applied during the application of Illumina/Ultima adapters. For Illumina sequencing, the indexing PCR was performed with the following recipe per well for a 96-well plate: (final volume is 50 µL): 21 µL eluted DNA (see above), 25 µL Q5U 2X Master Mix (New England Biolabs #M0597L), 2 µL 10 µM TruSeq i5 primers, 2 µL TruSeq i7 Flow Cell primers, and 0.5 µL EvaGreen 100X (Biotium #31019). For the Ultima libraries, indexing PCR was the same as above except that instead of the i7 Flow Cell primer, we used a barcoded i7 primer with Ultima’s sequencing handle on the 5’ side and instead of the barcoded i5 primers, we used a primer that added on Ultima’s i5 sequencing handle. The plate was mixed and placed on a qPCR with the following thermal conditions: 98°C initial denaturation for 30 seconds, 98°C for 30 seconds, 57°C annealing for 20 seconds, 72°C extension for 20 seconds, 72°C plate read for 10 seconds (these last 4 steps were cycled until exponential amplification was seen; roughly 8-10 cycles).

#### Library concentration and cleanup

After PCR, 10-30 µL of each well was pooled and concentrated using a Nucleospin® Gel and PCR Clean-Up kit (Macherey-Nagel #740609) according to manufacturer instructions. Two equal volumes of Macherey-Nagel™ Binding Buffer NTI (Fisher Scientific # NC0958675) was added to the library pool. The solution was mixed and pulled through column using a vacuum manifold. The column was then washed 2x with 700 uL NT3 wash buffer. The column was spun (11,000 xg, room temp, 1 min) to remove all liquid. DNA was eluted by incubating the column with 50 µL Buffer EB (Qiagen #19086), incubating 2 min at room temp, and spinning again (11,000 xg, room temp, 1 min). The concentrated library was further cleaned with a 0.9:1 SPRI-cleaned using Mag-Bind® TotalPure NGS beads (VWR # 75877-716). The product was quantified using a Qubit and TapeStation. An additional round of SPRI-cleaning was performed if a high adapter dimer content was observed on the TapeStation trace.

### Sequencing

Libraries were sequenced across multiple platforms as follows. Initial quality control sequencing was performed on an Illumina NextSeq instrument using either a P1 100 or P2 200-cycle kit. This data was used to produce target cell count and read depth estimates for larger-scale sequencing runs. The first run containing MECP2*^+/y^* rostral/caudal cortex, hippocampus, thalamus, and striatum data at all ages (sciMETv3 BSC+SL protocol) was sequenced on an Illumina NovaSeq S4 flowcell at the OHSU Massively Parallel Sequencing Shared Resource center. The second large batch was MECP2*^+/y^* rostral/caudal cortex, hippocampus, thalamus, and striatum data at all ages (sciMETv3 BSC+LA protocol) sequenced over five Ultima UG 100 wafers at Ultima Genomics (Fremont, CA). The fourth batch was a MECP2^+/y^ and MECP2^W104X*/y*^ pilot wafer across the P28 caudal cortex, hippocampus, thalamus, and striatum (sciMETv3 BSC+SL protocol) over one Ultima UG 100 wafer via Ultima Genomics (Fremont, CA). This pilot was used to prioritize the hippocampus and caudal cortex for a larger-scale fifth batch across the aforementioned regions and all ages sequenced over four Ultima UG 100 wafers (sciMETv3 BSC+LA protocol) through Novogene (Sacramento, CA). Data was rigorously monitored for any sequence platform-driven artifacts. All parameters are included in sample metadata.

### Initial processing of FASTQ files

Following sequencing, reads were first demultiplexed using unidex (github.com/adeylab/unidex) by matching to a whitelist of expected barcodes allowing a hamming distance of 2 for each of 3 specified indexes: 1 incorporated during PCR, 1 during ligation, and a third during tagmentation, which makes up the first 9 bp of read 2. Bases 9-29 of read 2 were then removed, as they contain the transposase mosaic end recognition sequence (ME). Reads were iteratively trimmed for all possible adapter sequences using cutadapt (v4.1).

Alignment was then performed to mm10 using BSBolt (v1.4.8) with the custom wrapper script fastq_align.pm at github.com/adeylab/premethyst. This script runs the aligner with reads 1 and 2 swapped due to the opposite configuration of sciMET adapters compared to traditional bisulfite sequencing adapters, followed with read sorting by name (cell barcode). mm10 was used for compatibility to annotated atlases.(*33*, *34*) PCR duplicates were then removed for each cell using Premethyst script bam_rmdup.pm and then methyl calls for CG and CH contexts were extracted using Premethyst script bam_extract.pm. All calls were then wrapped into an .h5 file using h5py (v3.15.1) in a custom Python script. The .h5 file is organized such that the first level contains groups named for each methylation context, the second level contains groups for each cell barcode, and the third contains datasets with corresponding base-call information for each captured cytosine along with aggregated counts. Lastly, aggregated 100,000 and 1500×500 window counts were calculated and stored in the .h5 files with the amethyst-facet (v1.2.4) *agg* command.

### Resolution and clustering of distinct populations

In R, an empty Amethyst (v1.0.4) object was built with *createObject().* Paths to h5 files were populated manually. Intermediate Premethyst metadata files generated during initial processing containing coverage and global methylation metrics were added using helper functions *addCellInfo* and *addAnnot.* To select for high-coverage cells, the metadata slot was filtered to cells containing at least 900k cytosines captured using *dplyr::filter* (v1.1.2). The rows corresponding to each chromosome for every cell were cataloged using the *indexChr* function. Following chromosome indexing, methylation values over 100kb windows were loaded into the object using the *loadWindows* command. Three genomeMatrices went into the final dimensionality reduction: 1) %mCH levels over 100kb windows with at least 10 values observed in at >80% of cells; 2) %mCH values over top 3,000 gene bodies (defined as the start to end base position in the mm10 Gencode annotation file) with the highest standard deviation between groups at P28; and 3) mCG scores over 100kb genomic windows using at least 5 values observed in >30% of cells. Score - a normalized measure of deviation from baseline - was calculated as: (mCG_feature_ - mCG_global_)/(1 - mCG_global_) if mCG_feature_ - mCG_global_ > 0, and (mCG_feature_ - mCG_global_)/(mCG_global_) if mCG_feature_ - mCG_global_ < 0.

After calculating methylation levels over genomic features, we calculated truncated singular values for each context using the Implicitly Restarted Lanczos Bidiagonalization Algorithm (IRLBA)(*139*) with *runIrlba*. To alleviate some sparsity-induced artifacts in the CG matrix, We also ran the *regressCovBias*(method = “gam”) function and *RunHarmony*(nclust = 20, kmeans_init_nstart = 1, kmeans_algorithm = “Lloyd”, sigma = 0.2, max_iter = 10) on the irlba result. We then combined the first 30 dimensions from CH percent over 100kb windows, 10 dimensions from CH percent over gene bodies, and 20 dimensions from CG score over 100kb windows into one multimodal reduced dimensionality result. This was used as input for *runCluster(k = 100, method = “louvain”)* and *runUmap(neighbors = 100,, dist = 0.1, method = “euclidean”).* All clustering methods can be found in the associated code for this manuscript, which will be posted on Github.

### Cell type annotation

Biological identity of the distinct groups was determined by consensus information from multiple approaches. The following approach was performed separately for each brain region and age. First, we leveraged two pre-annotated single-cell mouse methylation atlases(*33*, *34*) to compare methylation profiles directly. Publicly available data was downloaded and mean %mCH over gene bodies, %mCH over 100kb windows, and %mCG over 100kb windows for each subtype were calculated. We then performed a Pearson correlation analysis between each age, brain region, and cluster for values over shared regions. Top five correlations were compared and the cluster was annotated by the public data “type” classification if subtypes were in consensus. The canonical mCG context tended to produce higher correlations for P7 samples while the non-canonical mCH context tended to produce higher correlations for P14-P28 samples. Second, we visualized mCG patterns over canonical marker genes. Distinct hypo-mCG regions present across the gene body were used as a putative indicator of expression. Broad class annotation was validated in this manner - for example, by looking at patterns over *Slc17a7* to determine excitatory neuron populations. Third, mCH levels over gene bodies are known to be anticorrelated with expression. We used *aggregateMatrix* to calculate mean %mCH over gene bodies and compared this to trimmed mean data from the Allen Brain Institute(*140*) with the most negative correlation being a candidate for cell type assignment. Finally, we also took into account the dimensionality reductions to refine annotations, such as when correlation analysis produced two equally strong results, but one classification was clearly more likely based on neighbor annotation.

### Differentially methylated region analysis between successive ages

Counts from 1500 x 500 sliding windows previously calculated with facet *agg* -u 500=1500:500 were loaded into Amethyst with the *loadSmoothedWindows*(*nminValues = 10, nminCells = 2*) function by type and age. *testDMR(nminTotal = 10, nminGroup = 10)* was then run between each successive age within a subtype. *testDMR* is Amethyst’s core differentially methylated region testing function, which constructs contingency tables of c and t sums in member and nonmember cells, then performs a variation of a two-sided Fisher’s exact test from methylKit(*141*) to calculate a two-tailed *p*-value approximation using hypergeometric distribution probabilities with a continuity correction. Results were filtered using *filterDMR(method = “BH”, pThreshold = 0.05, logThreshold = 1.25, correctionLevel = “withinGroup”)* and collapsed with *collapseDMR(maxDist = 2000, minLength = 1500)*. The same parameters were used for CG and CH methylation. However, to keep analyses in **Fig. 3A-B** comparable, uncollapsed 500bp DMRs were used.

### Reactome Pathway analysis

Analysis in **Fig. 3C** was done using the ReactomePA (v1.14.0) R package. Genes with more than 5 significant 500bp CG DMR windows and 30 CH DMR windows within each major type were tested for enrichment separately. Entrez IDs were first extracted from gene symbols using the *bitr* function from clusterProfiler (v4.8.2) and mapping to org.Mm.eg.db (3.17.0). Enrichments were then tested for using the *enrichPathway* function with default settings. Results were then filtered for *p_adj_* < 0.05 and count > 2. In **Fig. 3C**, all significant enrichments are shown that were enriched in 5+ major types. To extract genes contributing to each enrichment, the *mapIDs* function was used.

### Low methylation region analysis

To identify CG LMRs in **Fig. 3B**, a sum matrix calculated from *loadSmoothedWindows* was generated using default settings. Windows with under 100 total c and t counts across all groups were removed. Groups that had <= 200 cells were not tested, nor were windows that had < 20 c and t observations within a group. After these filtering steps %mCG was calculated in passing windows. Windows that had <= 10% CG were considered LMRs. Specific LMRs were classified as windows that were identified in 1:3 groups. LMRs in (*n*_groups_ - 10): *n*_groups_ were considered shared, because many groups with low methylation in LMRs did not pass the strict filtering measures.

To identify CH LMRs in **Fig. 3C**, a sum matrix calculated from *loadSmoothedWindows* was generated using default settings. To account for low global %mCH levels at early postnatal time points, the sum matrix was filtered to only include P28 populations with > 2% global mCH, hence the discrepancy in max populations between **Fig. 3B-C**. Windows with under 100 total c and t counts across all groups were removed. Groups that had <= 200 cells were not tested, nor were windows that had < 20 c and t observations within a group. After these filtering steps %mCH was calculated in passing windows. Any window with <= 1% mCH was considered a CH LMR. Similar to CG testing, Specific LMRs were classified as windows that were identified in 1:3 groups. LMRs in (*n*_groups_ - 10): *n*_groups_ were considered shared, because many groups with low methylation in LMRs did not pass the strict filtering measures.

To test for overlap with genomic features, a list of candidate regulatory elements (cCREs) for mm10 was downloaded from ENCODE and converted to genomic ranges. Gene body locations were extracted from the GENCODE mm10 gtf annotation file for protein coding genes and the start and end were extended by 5kb. Exons were also extracted from the GENCODE annotation file and subtracted from genebodies with the *setdiff* function from GenomicRanges(v1.52.1) to get a list of introns and exons. The *reduce* function was used to generate non-overlapping segments of introns and exons. After acquisition of the feature lists, distinct LMRs were first intersected with cCREs using the *foverlaps(type = “any”, nomatch = NA)* function from data.table (v1.17.8). Any duplicate overlaps were removed. Then, regions that did not map to cCRE sites were intersected with the list of GENCODE introns and exons using the same strategy. The result was visualized using the ggplot2 (v4.0.2) *geom_col(position = “fill”)* function to show the proportion of LMRs overlapping each genomic element.

### mCH burst identification and analysis

mCH bursts were identified in the following manner. A sum matrix calculated from *loadSmoothedWindows* was generated using default settings. To account for low global %mCH levels at early postnatal time points, the sum matrix was filtered to only include P28 populations. Only groups with > 100 cells were included. For each major type, the sum matrix was filtered to include windows where observations were recorded in at least 75% of groups and the mean was calculated for passing windows. To reduce noise for slope detection, the matrix was smoothed with data.table’s *frollmean(n = 3, align = “center”)* function. Changepoints were then detected in the mean signal with the changepoint (v2.3) package’s *cpts* function using default settings. Changepoints within 4000 bp of each other were flagged as neighbors. To specifically identify spikes rather than depressions, neighboring changepoints were filtered where a positive slope preceded a negative slope. Finally, changepoint sets were filtered where the minimum value in the 10 surrounding windows was < 2% mCH. The *foverlaps* function was then used to detect neighboring protein coding genes (within 15kb) extracted from the GENCODE mm10 gtf annotation file.

To test for ontology enrichments of neighboring genes (**Fig. 4F**), the topGO (v2.52.0) package was used. Only recurrent (>6) overlaps were tested with the *GOdata(ontology = “BP”, nodeSize = 10)* function, followed by *runTest(algorithm = “elim”, statistic = “fisher).* As the elim algorithm inherently considers hierarchical relationships within the gene ontology structure and eliminates redundant testing, the developers consider the resulting *p*-values corrected.(*142*) Results were filtered for *p_elim_* < 0.01, significant contributing genes > 3, and fold change > 2. To reduce redundancy of GO terms, the *calculateSimMatrix(ont = “BP”, method = “Rel”)* and *reduceSimMatrix(threshold = 0.9)* functions from rrvgo (v1.12.2) were applied to results. The igraph (v2.1.4) package was then used to build a graph with the *graph_from_data_frame* function where edges were parent terms and weights are -log_2_(*p_elim_*). Finally, the circlepack plot in **Fig. 4F** was generated from this graph object using the *ggraph* function from the associated package (v2.2.1). Corresponding genes associated with cell fate specification terms were extracted using the topGO *genesInTerm* function.

To assess enrichment of mCH burst locations with ChIP seq data, the ReMap 2022 atlas was downloaded from the UCSC genome browser.(*109*) The complete atlas was filtered to data collected from brain-related biotypes along with blood for a negative control. Intersection with burst locations was calculated for each dataset using the *foverlaps* function. For each factor, enrichment was evaluated using a Fisher’s exact test by comparing the number of overlapping and non-overlapping regions in mCH bursts relative to the brain-related subset of ReMap peaks. Resulting *p* values were corrected for multiple testing with *p.adjust(method = “BH”)* from the stats (v4.3.0) package.

### Trajectory analysis for medial ganglionic eminence-derived interneurons

MECP2^+/y^ Sst Gaba, Pvalb Gaba, and young MGE Gaba populations were first subset from the final constructed object. The *irlba(dims = 2)* function from irlba (v2.3.5.1) was run on the pre-computed %mCH levels over 100kb windows observed in >80% cells. Irlba component 1 strongly reflected global mCH while component 2 reflected subtype differences. The *DiffusionMap(n_eigs = 2)* function from the destiny (v3.14.0) package was used to generate diffusion components from this dimensionality reduction. From the diffusion component result, the *slingshot* function from the Slingshot (v2.8.0) package was used to generate corresponding trajectory paths, and *DPT* from destiny was applied to determine endpoints. The *runCluster(k = 50)* function in Amethyst was then used to identify distinct populations, which were assigned to a Pvalb and Sst branch. Cells were divided into ten even bins for each branch according to rank along irlba component 1. For each bin, mean CG and CH levels were pseudobulked into 1500×500bp windows using the *loadSmoothedWindows* function under default settings. To determine diverged loci between branches, *testDMR(nminTotal = 10, nminGroup = 10)* was run between the top 2 Pvalb and Sst bins. Results were filtered with *filterDMR(method = “BH”, pThreshold = 0.05, logThreshold = 1.25)* and collapsed with *collapseDMR(maxDist = 2000, minLength = 3000)*.

### Trajectory analysis for intratelencephalic cortical glutamatergic neurons

MECP2^+/y^ young and mature IT CTX Glut populations were first subset from the final constructed object. The *irlba(dims = 10)* function from irlba (v2.3.5.1) was run on the pre-computed %mCH levels over 100kb windows observed in >80% cells. The *DiffusionMap(n_eigs = 3)* function from the destiny (v3.14.0) package was used to generate diffusion components from the first 5 components of this dimensionality reduction. The *runCluster(k = 50)* function in Amethyst was used to identify distinct populations from the first 10 irlba components, which were assigned to individual branches. Since young L4/5 IT CTX Glut cells mapped to both the L4 IT CTX Glut and the L5 IT CTX Glut branch, the population was duplicated and analysis continued with one set mapping to each branch respectively. From the diffusion component result, the *slingshot* function from the Slingshot (v2.8.0) package was used to generate corresponding trajectory paths with specified start and end clusters. Cells were divided into ten even bins for each branch according to rank along diffusion mapping component 1. For each bin, mean CG and CH levels were pseudobulked into 1500×500bp windows using the *loadSmoothedWindows* function under default settings. To determine diverged loci between branches, *testDMR(nminTotal = 10, nminGroup = 10)* was run between the top 2 bins in the branch of interest vs. top 2 bins in each of the 3 other branches. Results were filtered with *filterDMR(method = “BH”, pThreshold = 0.05, logThreshold = 1.25)* and collapsed with *collapseDMR(maxDist = 2000, minLength = 3000).* To hierarchically cluster CG and CH loci in **Fig. 5F-I**, mean %mC values for each DMR locus were calculated using the *amethyst::makeWindows(nmin = 5)* function. Mean values for each branch and pseudotime bin were calculated using *amethyst::aggregateMatrix(minValues = 5).* the *pheatmap* function from pheatmap (v1.0.13) was used to hierarchically cluster the rows (DMR loci) into 10 distinct groups for CG methylation and 20 distinct groups for CH methylation using default settings. In **Fig. 5F,H**, values only up to the top 25 loci per cluster are shown.

Differentially methylated region and global analysis for MECP2^+/y^ vs MECP2^W104X/y^ cells

Differentially methylated regions (DMRs) were identified with Amethyst (v1.0.4) in the following manner for MECP2^+/y^ vs MECP2^W104X/y^ cells. First, the sum of MECP2^+/y^ and MECP2^W104X/*y*^ c (methylated) and t (unmethylated) observations were pseudobulked by brain region, sequencing date, type, genotype, and mouse in a 1500bp x 500bp sliding window using the *loadSmoothedWindows(nminValues = 10)* function. Windows failing to meet the threshold of 10 c and t observations recorded in either member or non-member groups were removed. For the remaining contingency tables of c and t sums in member and nonmember cells, a variation of a two-sided Fisher’s exact test was used(*141*) to calculate a two-tailed *p*-value approximation using hypergeometric distribution probabilities with a continuity correction with *testDMR*. *testDMR* is Amethyst’s core differentially methylated region testing function, which constructs contingency tables of c and t sums in member and nonmember cells, then performs a variation of a two-sided Fisher’s exact test from methylKit(*141*) to calculate a two-tailed *p*-value approximation using hypergeometric distribution probabilities with a continuity correction. Filtering was then done with *filterDMR(method = “BH”, pThreshold = 0.1, logThreshold = 1(CG)* or *0.5(CH), correctionLevel = “withinGroup”)*. To account for any batch or sequencing-induced effects, two comparisons were performed. The first was between littermate (or as close as possible) matches processed and sequenced in the same day. The second was between all other replicates. Only significant results with a mean *p*_adj_ < 0.05 and mean log_2_FC > 1 were kept. We consider this consensus-based approach quite stringent given that multiple testing correction is applied for millions of windows tested. To keep results in **Fig. 6E** comparable, values shown indicate the number of significant differentially methylated 500bp windows, not collapsed DMRs. For global results in **Fig. 6D**, mean mCG and mCH values were calculated per brain region, sequencing date, type, genotype, and mouse. Mean global MECP2^W104X/y^ mC were normalized to mean global MECP2^+/y^ mC values processed in the same prep and sequencing batch to account for any technical discrepancies. A two-sided t test was then performed on these mean ratios per biological replicate to test for shift in distribution from *mu = 1*. The resulting *p* values were corrected for multiple testing at the level of each age and subtype using *p.adjust(method = “BH”)* from the stats (v4.3.0) package.

## Acknowledgments

The authors would like to thank Dr. Mati Nemera and Dr. Harrison Gabel for their contributions to analyses of global mCH levels in parvalbumin-positive interneurons following MECP2 loss. We are also grateful to Dr. Longzhi Tan for generously providing MALBAC-DT metadata associated with PMID:33484631 and for writing an exceptionally thorough methods section. The authors would like to recognize Dr. Sonia Acharya for her contributions to the development of the neuron staining and flow sorting protocol, and Andrew Fields for regular assistance with the Sony SH800 flow sorting instrument and reagent acquisition. Finally, we thank members of the Adey, Saunders, and O’Roak labs at Oregon Health & Science University for their valuable feedback and discussion throughout this work.

## Funding

Medical Research Foundation Early Clinical Investigator Grant (LER)

National Institutes of Health BRAIN Initiative RF1MH128842 (ACA)

Silver Family Foundation Innovator Award (ACA)

National Institutes of Health NINDS R01NS110868 (GM)

## Author contributions

Conceptualization: LER, ACA, GM

Methodology: LER, RVO, BLO, SK, JFY, GM, AS, ACA

Investigation: LER, RVO, BLO, SK, JFY, GM, AS, ACA

Visualization: LER, ACA

Funding acquisition: LER, ACA, GM

Project administration: LER, SK, JFY, ACA, GM Supervision: LER, ACA, AS, GM

Writing – original draft: LER

Writing – review & editing: LER, ACA, GM

## Data, code, and materials availability

All raw and processed data files will be made publicly available on the National Center for Biotechnology Information Gene Expression Omnibus (NCBI GEO) website. Initial processing of sequencing output to base-level methylation calls can be performed with Premethyst commands, which are available at github.com/adeylab/premethyst. Amethyst is publicly available for installation at github.com/lrylaarsdam/amethyst. All code used for analysis of this dataset will be made available on GitHub and Zenodo.

